# Temporal change in chromatin accessibility predicts regulators of nodulation in *Medicago truncatula*

**DOI:** 10.1101/2021.08.07.455463

**Authors:** Sara A. Knaack, Daniel Conde, Sanhita Chakraborty, Kelly M. Balmant, Thomas B. Irving, Lucas Gontijo Silva Maia, Paolo M. Triozzi, Christopher Dervinis, Wendell J. Pereira, Junko Maeda, Henry W. Schmidt, Jean-Michel Ané, Matias Kirst, Sushmita Roy

## Abstract

Rhizobia can establish symbiotic associations with legumes to provide plants with nitrogen needed in agricultural systems. Symbiosis triggers extensive genome and transcriptome remodeling in the plant, yet the extent of chromatin changes and impact on gene expression is unknown. We profiled the temporal chromatin accessibility (ATAC-seq) and transcriptome (RNA-seq) dynamics of *M. truncatula* roots treated with rhizobia lipo-chitooligosaccharides. Using a novel approach, Dynamic Regulatory Module Networks, we predicted gene expression as a function of chromatin accessibility and accessible *cis*-regulatory elements. This approach identified the *cis*-regulatory elements and associated transcription factors that most significantly contribute to transcriptomic changes triggered by lipo-chitooligosaccharides. Regulators involved in auxin (IAA4-5,SHY2), ethylene (EIN3, ERF1) and abscisic acid (ABI5) hormone response, as well as histone and DNA methylation (IBM1), emerged among those most predictive of transcriptome dynamics. RNAi-based knockdown of EIN3 and ERF1 reduced nodule number in M. truncatula validating the role of these predicted regulators in symbiosis between legumes and rhizobia.

**Significance Statement:** Legumes can fix nitrogen through symbiosis with rhizobia in root nodules, a critical mutualistic relationship for crop productivity and agricultural sustainability. Introducing this symbiotic relationship into non-legume crops is of great interest, but limited knowledge of host genome modifications induced by rhizobia has hampered such efforts. We applied time-course analysis of chromatin accessibility and gene expression of *M. truncatula* roots treated with rhizobia lipochitooligosaccharides. We show that extensive remodeling of genome accessibility drives a large component of the temporal transcriptome dynamics. By predicting gene expression as a function of accessibility of regulatory features, we identified known and novel regulators that are associated with early nodule development, which may be critical for its engineering into crops.

## Introduction

Legumes can establish a well-characterized mutualism with nitrogen-fixing rhizobia. Signal exchanges between the host plant and bacteria initiate intracellular infection of host cells, followed by the development and colonization of root nodules (1). Nodules provide a unique niche for the bacteria and fix nitrogen. Nodulating plants can grow with little to no outside sources of nitrogen and even build soil nitrogen levels for subsequent crops (2). Hence, understanding symbiotic processes between legumes and rhizobia is extremely valuable for the productivity and sustainability of agricultural systems worldwide.

Symbiosis begins with compatible rhizobia detecting flavonoids and isoflavonoids produced by the legume host (3), and subsequent release of lipo-chitooligosaccharides (LCOs) by the bacteria. The host plant perceives LCOs with LysM domain receptor-like kinases heterodimers, such as Nod Factor Perception (NFP) and LysM Domain Receptor-like Kinase 3 (LYK3) in *Medicago truncatula* (4, 5). LCO perception activates a signaling cascade, involving the plasma membrane-localized LRR-type receptor kinase Doesn’t Make Infections 2/Nodulation Receptor Kinase (MtDMI2/MtNORK), the calcium-regulated calcium channel (MtDMI1), cyclic nucleotide-gated calcium channels, *M. truncatula* calcium ATPase 8 (MtMCA8), and including the components of the nuclear pore complex (6–8). The cascade results in oscillations of nuclear calcium concentrations, detectable by the nucleus-localized Calcium/Calmodulin-dependent protein Kinase (CCaMK, MtDMI3 in *M. truncatula*) (9). CCaMK activates the transcription factor (TF) Interacting Protein of DMI3 (MtIPD3/CYCLOPS). Downstream, other TFs are activated, such as Nodulation-Signaling Pathway 1 and 2 (NSP1 and NSP2), Nodule INception (NIN), Ethylene Response Factor required for Nodulation 1,2 and 3 (ERN1, 2 and 3), and Nuclear Factor YA-1 and YB-1 (NF-YA1 and NF-YB-1) (10, 11).

The coordinated activity of these TFs triggers transcriptional changes (12) essential for infection of the root hair cells (in *M. truncatula*), nodule organogenesis and infection of the nodule cortex (10, 11). These processes require changes in chromatin accessibility (13), i.e., the distribution and organization of nucleosomes. Densely packed nucleosomes limit access of regulators to genomic regions, yet nucleosomes are depleted at active regulatory loci. Nucleosome occupancy is dynamic and creates a continuum of accessibility from closed to open, which is important for cell function (14). In plants, chromatin reorganization regulates photomorphogenesis and flowering (15, 16). Active DNA demethylation by DEMETER (DME) is critical for gene expression reprogramming during nodule differentiation in *M. truncatula*, and the acquisition of organ identity (13). Also, in *M. truncatula*, the gene expression level of Nodule-specific Cysteine-Rich genes (NCR) across root nodule zones are correlated with chromatin accessibility (17).

The extent of chromatin accessibility change and impact on transcriptional regulation in rhizobial infection, colonization, and nodule development, remains unknown. Thus, we measured temporal changes in the transcriptome (RNA-seq) and genome-wide chromatin accessibility (ATAC-seq) in response to *Sinorhizobium meliloti* LCOs in *M. truncatula* roots. To characterize the gene regulatory network, the role of chromatin accessibility (and consequent impact on the transcriptional dynamics) we applied a novel algorithm, Dynamic Regulatory Module Networks (DRMN) (18), to predict gene expression as a function of chromatin accessibility profiles of *cis*-regulatory features. DRMN results suggest that chromatin accessibility and specific regulatory features (and associated TFs) play a critical role in regulating the transcriptional dynamics in response to LCOs.

## Results

### Root transcriptome response to LCOs involves genes activated by rhizobia and early nodule development in *Medicago*

We profiled the global transcriptomic changes of rhizobium LCO signaling with RNA-seq in *M. truncatula* using the Jemalong A17 genotype, treated with LCOs purified from *S. meliloti*. An LCO concentration of 10^−8^ M was used, as in previous studies (19, 20). Samples were analyzed for control (t=0 hr) and seven time-point conditions after treatment (15 and 30 min; 1, 2, 4, 8, and 24 hr). Principal component analysis (PCA) showed clustering of biological replicates and time-dependent ordering, the first component explaining ∼36% of variation (**Fig. 1A, S1**). Comparison of expression levels at each time point (relative to control, t=0 hr) revealed 12,839 differentially expressed (DE) genes with significant change in expression (adjusted-*P* < 0.05), including 7,540 and 7,051 up-and down-regulated at one time point relative to control, respectively.

**Figure 1.**
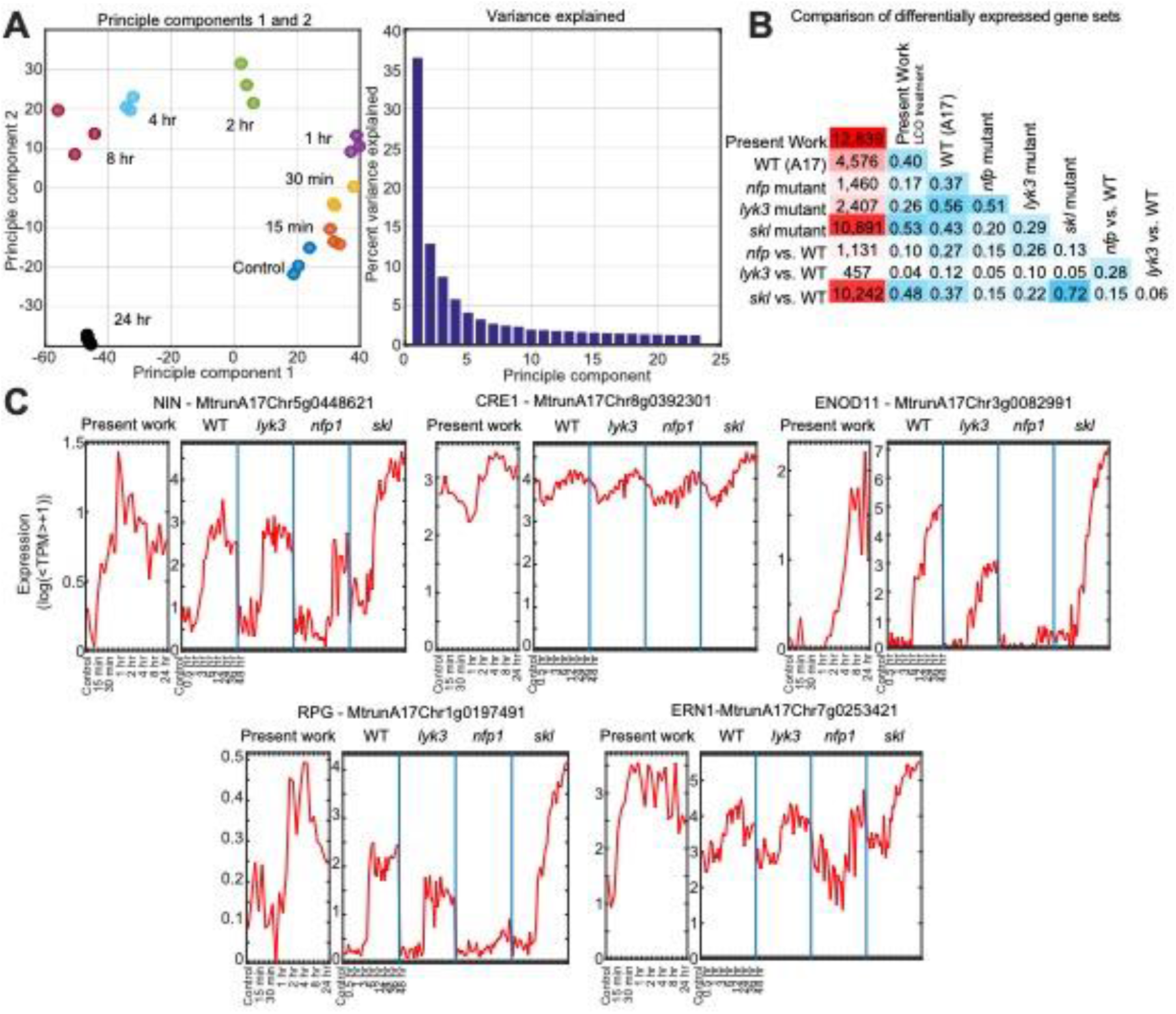
Transcriptome data summary. **(A)** Principal components analysis (PCA) showing grouping and ordering of biological replicates across time points. Principle components 1, 2, and 3 explain ∼50% of temporal variation. **(B)** Similarity scores (F-score) between the DEG set obtained in this study (LCO treatment) and DEG sets identified from previously published time-course data under rhizobium treatment from Larrainzar et al. (12). For the latter data, DE genes were called with respect to control for each respective time course (rows and columns corresponding to WT, *nfp, lyk3, skl*), and with respect to WT at each time-point for each mutant strain (rows and columns with “vs. WT” labels). **(C)** Expression patterns of *NIN, CRE1, ENOD11, RPG* and *ERN1* across time points in this study, and four genotypes analyzed previously (12).

To corroborate these results with previous work on transcriptome dynamics of symbiosis, the identified DEGs were compared to DEGs identified from a published time course data of *M. truncatula* roots inoculated with rhizobium from Larrainzar et al. (12) (**Fig. S2A**). Comparisons were made to DEGs in the following genotypes: Jemalong A17 wild type, LCO-insensitive *nfp* mutant, infection *lyk3* mutant, LCO-hypersensitive *skl* mutant. The highest similarity, measured by F-score, to our DEG set was for the mutant genotype most sensitive to LCOs, *skl* (0.53), and the wild-type (WT) strain (0.40). Marker genes for rhizobium-induced nodulation were up-regulated (compared to t=0 hr), including *NIN* (*Nodule inception*, induced after 15 min, with a maximum induction at t=1 hr), *CRE1* (*Cytokinin Response 1*, at 4 hr, 8 hr, and 24 hr), *ENOD11* (*Early nodulin 11*, highly induced at 8 and 24 hr), *RPG* (*Rhizobium-directed Polar Growth*, at 4 hr) and *ERN1* (*Ethylene Responsive Factor Required for Nodulation 1*, induced as early as t=15 min, **Fig. 1C**). The similarity was lowest for the *lyk3* (0.26) and *nfp* (0.17) mutants (**Fig. 1B, Fig. S2B** and **C**). While our DEGs had the greatest overlap with the *skl* genotype DEGs, we detected more DEGs compared to Larrainzar et al, likely due to differences in growth conditions (aeroponics versus agar plates) and treatment (purified LCOs versus *Sinorhizobium medicae*), both inducing a strong LCO response.

To examine more complex transcriptome dynamics beyond pairwise DE analysis associated with LCO response, we applied ESCAROLE, a probabilistic clustering algorithm designed for non-stationary time series (21). The expression data were clustered into seven modules at each time point (very low, low, medium-low, medium, medium-high, high and very high expression, **Fig. 2A**). Seven modules maximized the log-likelihood and silhouette index (**Fig. S3A, B**). Next, 12,261 transitioning genes (those changing module assignment over time) were identified, including several implicated in symbiosis (**Fig. S3C**). Transitioning genes with similar dynamics were clustered using hierarchical clustering, identifying 112 clusters (>=10 genes each) (**Fig. 2B**) including 11,612 genes (**Supplementary methods**). Among clusters representing downregulation of expression over time, several were enriched for Gene Ontology (GO) processes implicated in defense responses to bacterium (cluster 293, downregulated from 2-4 hr), and the biosynthesis of plant hormones involved in the suppression of nodulation (**Fig. 2C**). For instance, cluster 299 (downregulated after 2 hr) is enriched for jasmonic acid (JA) biosynthesis and JA response genes, including *Coronatine insensitive 1* (*COI1*), which forms part of the JA co-receptor complex for the perception of the JA-signal (22). Among the gene clusters upregulated over time, several are implicated in early stages of symbiosis and nodule development. For instance, cluster 186 (induced 2-4 hr after LCO treatment; **Fig. 2C**) is enriched in genes implicated in the regulation of meristem growth, including an *Arabidopsis trithorax 3* (*ATX3*) homolog (MtrunA17Chr4g0005621) and a *Lateral Organ Boundaries Domain* (*LBD*) transcription factor (MtrunA17Chr4g0043421). *ATX3* encodes an H3K4 methyltransferase (23), and LBD proteins are characterized by a conserved Lateral Organ Boundaries (LOB) domain and are critical regulators of plant organ development (24), including lateral roots and nodules (25). This cluster also contains *EPP1* and the cytokinin receptor *CRE1*, both positive regulators of early nodule symbiosis and development (26, 27). Other essential regulators of LCO signaling are also found in clusters exhibiting induction under LCO treatment (**Fig. S3D**), such as *DMI1* (cluster 197, **Fig. 2C**), *NIN* (cluster 205), *NF-YA1* (cluster 177) and the marker of LCO perception *ENOD11* (cluster 296). Together the DE and ESCAROLE analysis showed that *M. truncatula* response to LCOs is characterized by complex expression dynamics recapitulating several known molecular features of this process.

**Figure 2.**
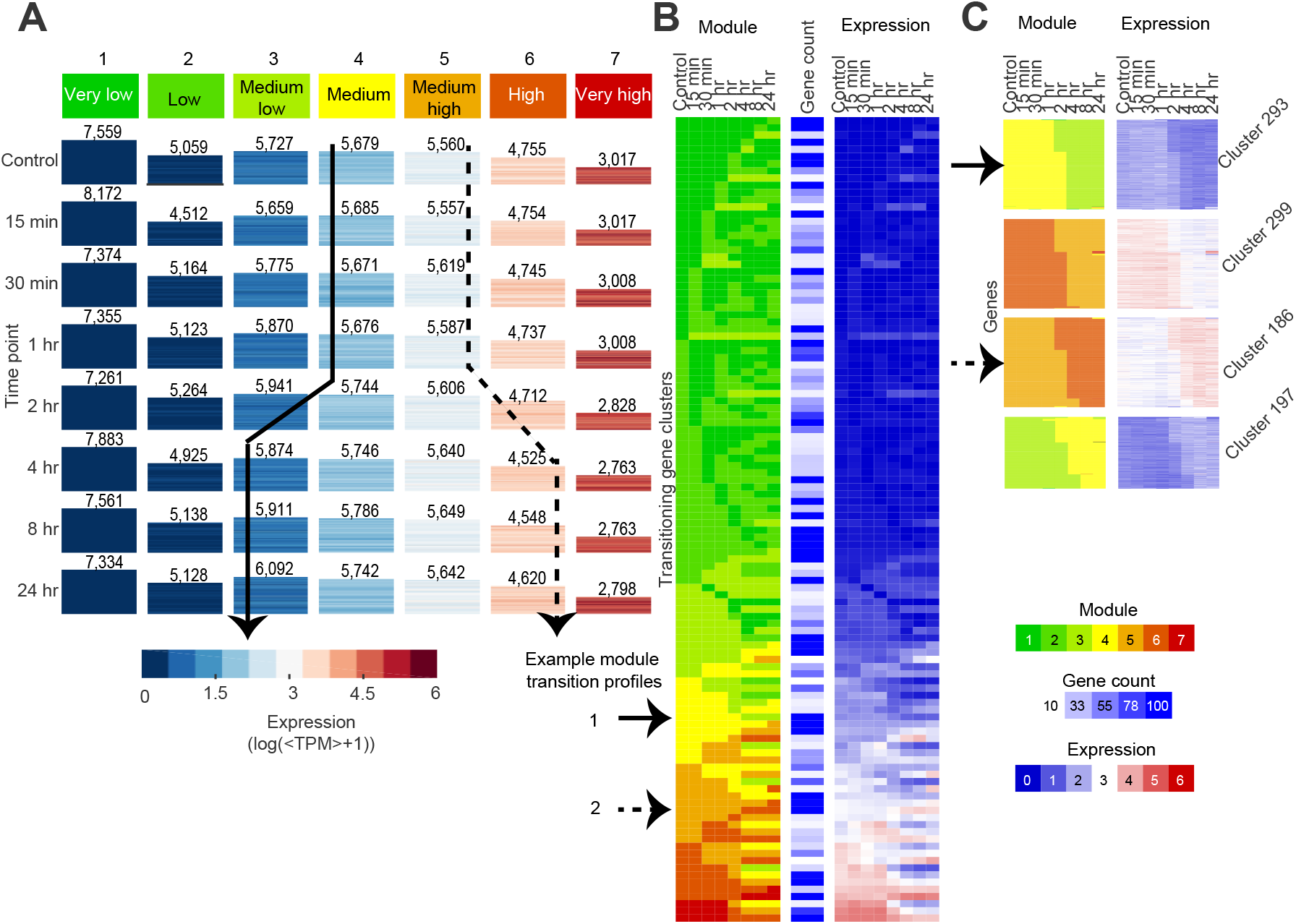
Transcriptome dynamics in response to LCOs. **(A)** ESCAROLE results for seven modules, based on transcript abundance data. Each heatmap includes genes assigned to that module at that time point, and the height of each corresponds to the number of genes (inset numbers). **(B)** The module assignment heatmap depicting typical gene expression trends obtained by hierarchical clustering of gene module profiles into transitioning gene sets. Shown are the mean module assignments, number of genes in each set, and expression levels at each time point for each cluster. Arrows indicate two example trends of expression change. **(C)** Examples of transitioning gene clusters showing gene expression upregulation or downregulation, enriched for genes implicated in nodulation such as defense response to bacterium (Cluster 293) or meristem growth (Cluster 186).

### LCO treatment causes genome-wide changes in chromatin accessibility

To study chromatin accessibility changes in a genome-wide manner in response to LCOs, we performed ATAC-seq on samples at all time points matching our RNA-seq time course. Overall, 54-235 million paired-end reads were obtained for each sample, with 46-75% mappable to the (v5) reference genome (**Fig. S4**). We evaluated chromatin accessibility in gene promoter regions, defined as ±2 kbp around the transcription start site (TSS). To quantify promoter accessibility we obtained the mean per base pair (per-bp) read coverage within each region, for each time point. For each time-point the log-ratio of per-bp read coverage in each promoter was taken relative to the global mean of per-bp coverage, quantile normalized across time points, followed by k-means clustering and PCA (**Fig. S5A)**. We partitioned the resulting 51,007 gene promoter accessibility profiles into six characteristic patterns (clusters) using k-means clustering (**Fig. 3A, Fig. S5B**). Clusters 1 (14,338 genes) and 6 (13,083 genes) exhibit general patterns of decrease and increase in accessibility, respectively, whereas clusters 2-5 (5,460-6,377 genes) present more transient variation. The correlation of accessibility between time points suggests an overall reorganization of promoter accessibility 1-2 hr after the treatment (**Fig. S5C**). The temporal change in accessibility is evident for the promoters of several nodulation genes, including *CRE1, CYCLOPS*, and *EIN2* (**Fig. 3B, Fig. S5D** and **E**). PCA of the promoter signals showed time-dependent variation (**Fig. 3C, Fig. S5A**), with the first component explaining >50% of the variance.

**Figure 3.**
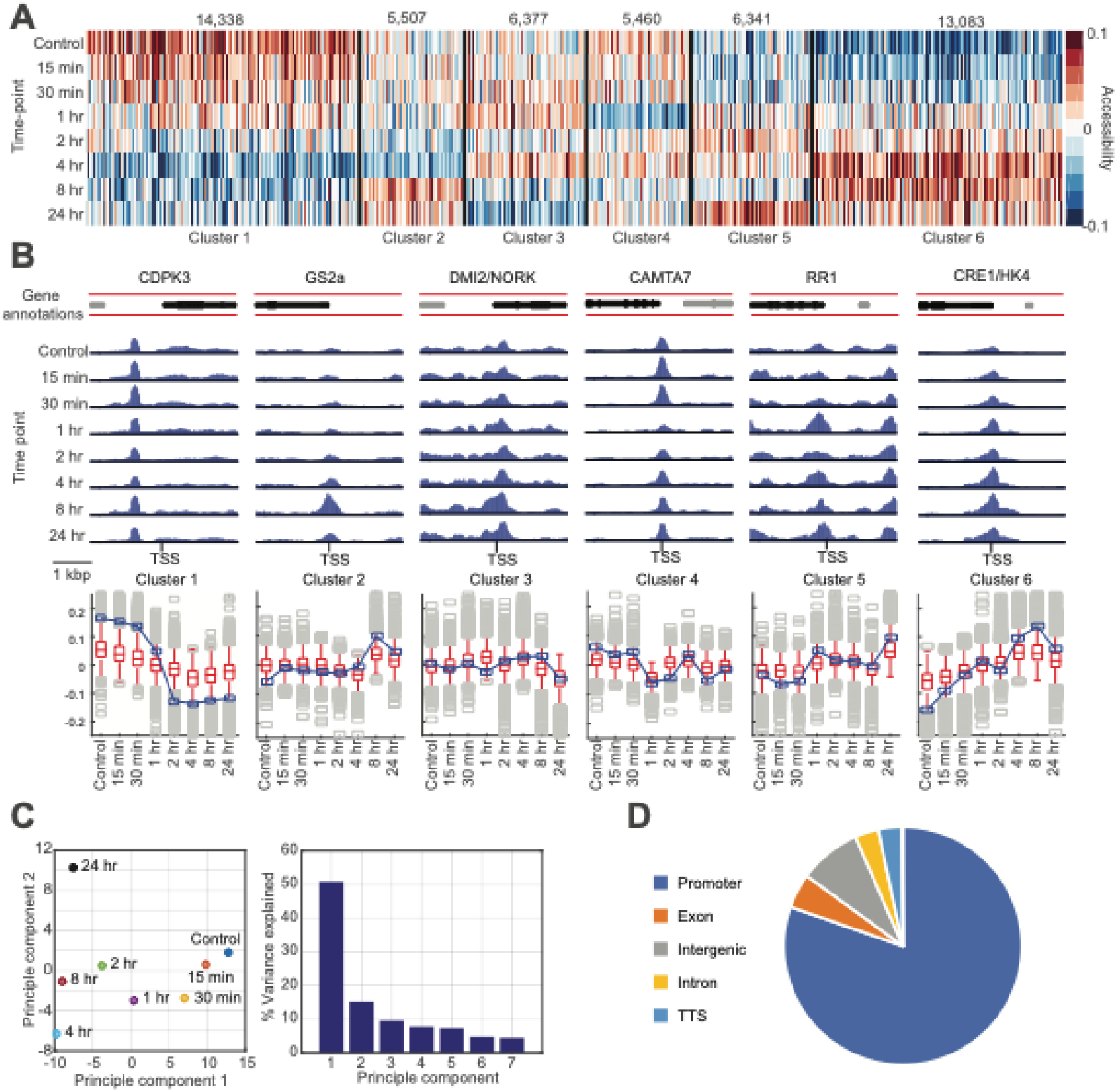
Chromatin accessibility data exploratory analysis. **(A)** Clustering of promoter accessibility profiles in the LCO-response time course. **(B)** Integrative Genomics Viewer (IGV) (29) plots of the normalized coverage profiles for the promoter regions (± 2 kbp from TSS) of genes involved in root nodulation, representing each cluster (upper panel). Gene annotation track (top) denotes the gene of interest (black), and neighboring genes (gray). **(C)** PCA results for the same promoter accessibility data. **(D)** Genomic annotations of the identified universal accessibility (ATAC-seq) peaks.

We called peaks for each time point using the Model-based Analysis of ChIP-Seq version 2 (MACS2) algorithm (28) (**Fig. S6A**), and merged peaks across time points with at least 90% overlap into *universal* peaks (**Fig. S6B**). Chromatin accessibility peaks showed a similar genomic distribution across time points, with 81-85% of peaks located within 10 kbp upstream and 1 kbp downstream of a gene TSS (**Fig 3D, S6C**) and spanning 50.4 Mbp (11.7%) of the *M. truncatula* (v5) genome. As with the promoter accessibility, clustering accessibility profiles of universal peaks identified distinct patterns of temporal change (**Fig. S6D** and **E**). Several of the clusters were associated with known TF motifs (**Fig. S6F**) and specific types of genomic regions (e.g. cluster 1 and 2) had higher proportions of intergenic peaks (**Fig. S6G**). These results suggest that LCO treatment had a genome-wide impact on chromatin accessibility, prospectively associated with simultaneous change in gene expression and regulation.

### Chromatin accessibility is correlated with transcriptional dynamics of nodulation genes

We evaluated the relationship between gene expression and chromatin accessibility at promoters (±2 kbp around the TSS) and universal peaks within 10 kbp upstream and 1 kbp downstream of a gene TSS. Correlating promoter accessibility and gene expression profiles identified 6,429 genes with significant correlation (**Fig. 4A**, *P*<0.05); 4,777 with positive correlation and 1,652 with negative correlation (**Fig. 4B**), representing 17.2% of the 37,356 genes analyzed. Among these were 36 genes with known roles in symbiosis (**Fig. S5D**), including *ERN1, CRE1, LYK10/EPR3, SKL/EIN2*, and *IDP3*/*CYCLOPS* with positive correlation, and *LYK8, ERN2, CAMTA3*, and *CAMTA4* with negative correlation. We next examined significantly correlated genes (**Fig. 4A**) and visualized those expression and accessibility profiles as ordered by the promoter accessibility clusters (**Fig. 3A**), separately for positive and negative correlation (**Fig 4B**). This revealed robust patterns of consistency between promoter accessibility and expression.

**Figure 4.**
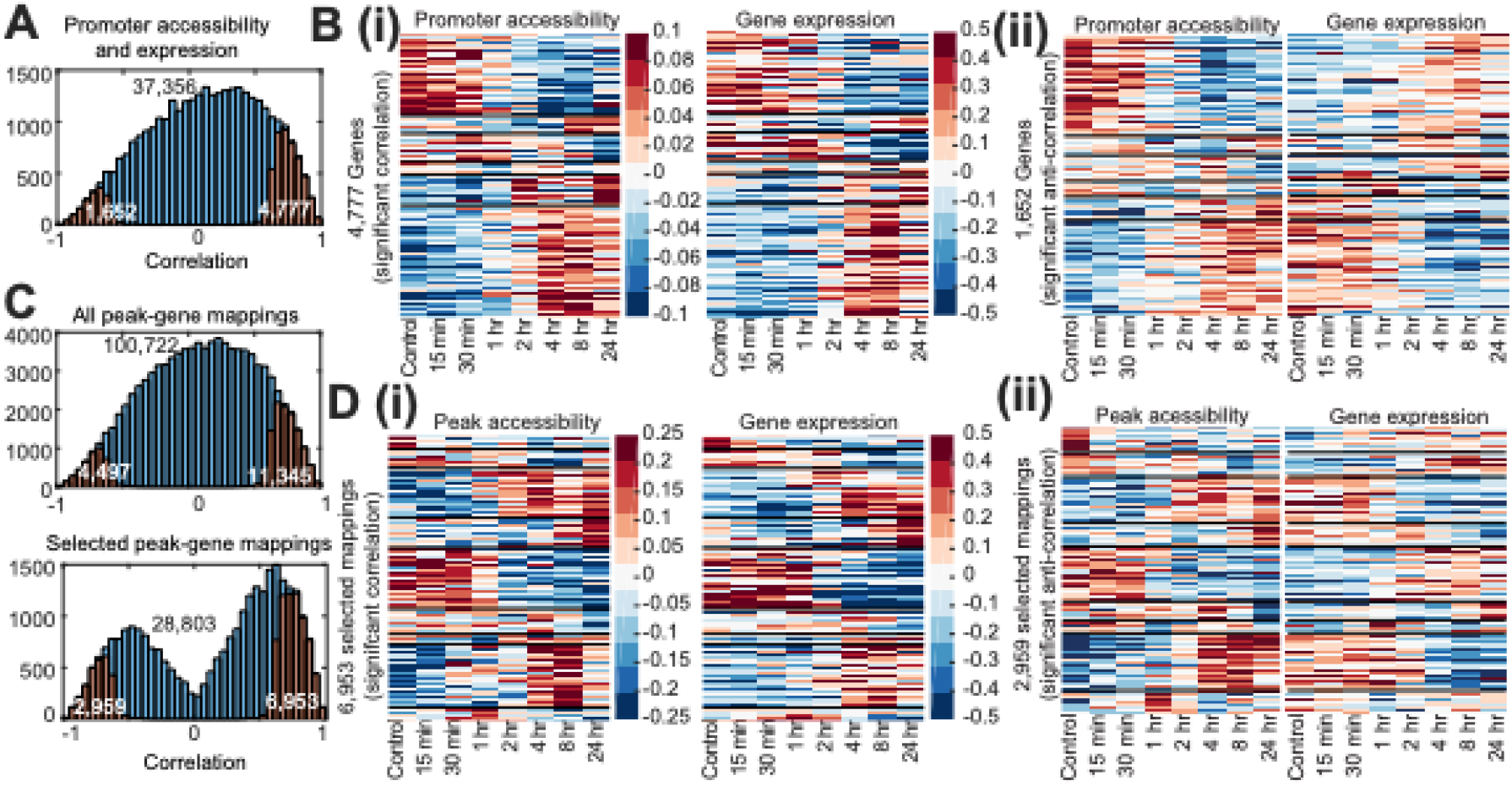
Correlation between chromatin accessibility and gene expression. **(A)** Histogram of Pearson correlation of all (blue) and significantly correlated (orange) promoter accessibility and gene expression pairs. The numbers are indicated with inset numbers. **(B)** Clusters of promoter accessibility and gene expression for significant (*P*<0.05) (i) positive and (ii) negative correlation relative to random. **(C)** Histograms of correlation for all (blue) and significantly correlated peaks and genes (orange), and associated statistics. The upper histogram includes all mapped peak-gene pairs, while the lower includes only the maximally correlated peak for each gene (below). **(D)** Clustered peak accessibility and corresponding expression profiles for significantly positively (i) or negatively (ii) correlated gene-peak mappings.

Correlating accessibility of universal peaks centered within 10 kbp upstream to 1 kbp downstream of gene TSSs identified 100,722 peak-gene mappings (out of a total 125,140) associated with 28,803 (of 37,536) expressed genes (**Fig. 4C, Fig. S6C** and **G**). Peak accessibility was significantly correlated with gene expression in 15.7% of these pairings (**Fig. 4C**), comparable to the 17.2% (6,429) genes with significant correlation between expression and promoter accessibility. When considering each gene and only the most correlated peak (28,803 selected pairs), 34.4% (9,912 genes) were significantly correlated, including 56 nodulation genes (**Fig. 4D**). Of these 9,912 genes presenting significant correlation, 5,735 (57.9%) do not present significant correlation with the corresponding promoter accessibility, indicating a prominent role for distal regulation (>2 kbp of gene TSS) for these genes.

Finally, the ESCAROLE-defined transitioning gene clusters exhibited coordinated trends between promoter accessibility and gene expression (**Fig. 2B, Fig. S3D**). 2,501 of the 11,612 (21.5%) transitioning genes exhibited significant correlation between their profiles of expression and promoter chromatin accessibility. These results suggest that chromatin accessibility is an important regulatory mechanism in transcriptional response to LCOs.

### DRMN integration of ATAC-seq and RNA-seq data identifies key regulators that determine gene expression dynamics in response to LCOs

To better understand how chromatin accessibility contributes to transcriptional changes in rhizobia-plant symbiosis, we applied Dynamic Regulatory Module Networks (DRMN) (18) to integrate the RNA-seq and ATAC-seq time course data. DRMN extends the ESCAROLE analysis (which examined only the transcriptome) by modeling the relationship between variation in accessibility and gene expression. DRMN predicts gene expression as a function of regulatory features (30) by first grouping genes into modules based on expression levels (similar to ESCAROLE), and then learning a regulatory program for each module. DRMN uses regularized regression and multi-task learning to incorporate the temporal nature of a data set (31) to simultaneously learn regression models for each module in each time point.

We applied DRMN with seven expression modules using two types of features (**Fig. 5A**): (1) the aggregated signal of ATAC-seq reads in gene promoters (± 2 kbp of the TSS), and (2) the ATAC-seq signal in genomic coordinates of known motifs within -10 kbp and +1 kbp of the TSS. Both feature types represent chromatin accessibility, but the first is independent of the presence of known motifs, whereas the second captures the accessibility of motif sites. Motif features were based on the CisBP v1.2 database for *M. truncatula* (32), and curated motifs of several known regulators of root nodulation, including CYCLOPS, NSP1, NIN and the nitrate response *cis*-element (NRE). Hyper-parameters for DRMN were selected using a grid search and quality of inferred modules (**Fig. S7A**). The DRMN modules represent statistically different expression levels (**Fig. S7B**, Kolmogorov-Smirnov test *P* <10^−300^). To assess the extent to which DRMN captures variation in expression, we correlated predicted and measured expression levels (**Fig. 5B, Fig. S7A, C**). The mean Pearson correlation of predicted and measured values per module was 0.26-0.46 (**Fig. S7C**) across all modules and time points, the least expressed module being most difficult to predict. (**Fig. 5C**) is also indicated. Comparing the genes in each module showed that the modules are more similar (F-score 0.88-0.94) before and after 2 hr, than across this time point (F-score<0.80), suggesting a significant module reorganization at ∼2 hr. This is consistent with the general reorganization of promoter accessibility ∼1-2 hr after the treatment (**Fig. S5C**). We additionally tested the modules for enrichment of known motifs (**Fig. S8**) and Gene Ontology (GO) processes (**Fig. S9**). Several regulators (e.g., KNOX and EDN transcription factor family members) and processes relevant to symbiosis were identified, including nodule morphogenesis, root-hair elongation and the MAPK cascade, as well as others relating to gene regulation and chromatin organization.

**Figure 5.**
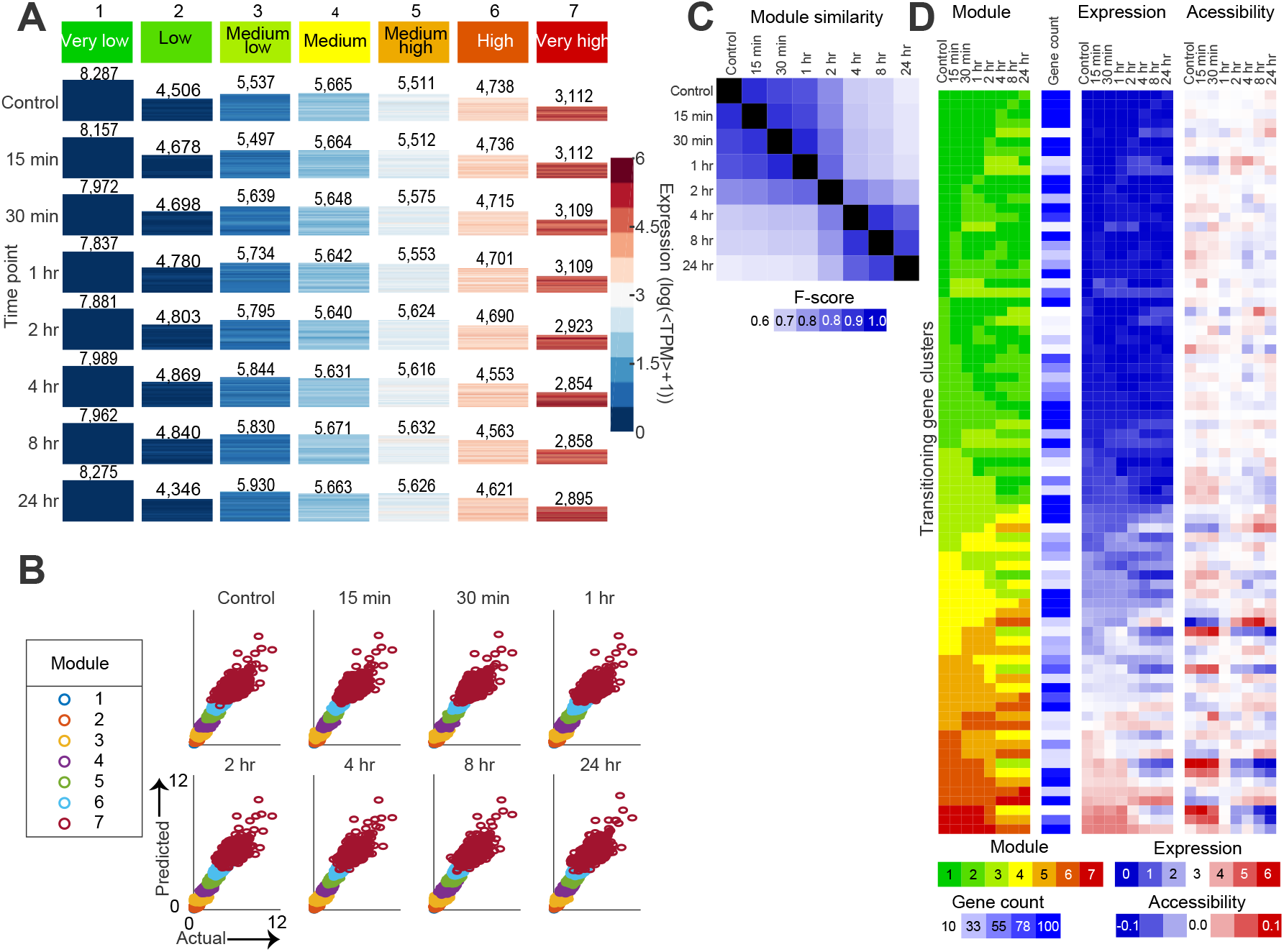
Dynamic Regulatory Module Network analysis. **(A)** Heatmap of DRMN inferred expression modules across the time course. Each heatmap corresponds to an expression module for each timepoint, the size of the heatmap corresponding to the number of genes assigned to that module (listed on top). **(B)** Scatter plots of actual and predicted values. **(C)** F-score similarity of DRMN modules across time points. **(D)** Shown are the mean DRMN module assignment, number of genes, expression levels and mean promoter accessibility levels for each inferred cluster (rows) across time (columns).

We used the DRMN results to prioritize regulators that shape transcriptional response to LCOs. Specifically, we identify regulators whose coefficient changed significantly (T-test *P*<0.05) between 0-2 and 4-24 hr, corresponding to the reorganization of expression modules (**Fig. 5C**). According to this criterion chromatin accessibility of gene promoters was an important predictor of expression for highly expressed genes (“Promoter ATAC-seq” for modules 5 and 6, **Fig. 6A**). We also identified the TFs IBM1 (Increase in BONSAI Methylation 1), ERF1 (Ethylene Response Factor 1), EDN1-3 (ERF Differentially Regulated During Nodulation 1, 2 and 3), EIN3 (Ethylene Insensitive 3), SHY2 (Short Hypocotyl 2), ABI4-5 (Abscisic Acid-Insensitive 4 and 5), MTF1 (MAD-box Transcription Factor 1), MtRRB15 (type-B response regulator 15), as well as several markers of meristem cells, KNOX and PLT (PLETHORA) protein families as important regulators (**Fig. 6B, Fig. S8**).

**Figure 6.**
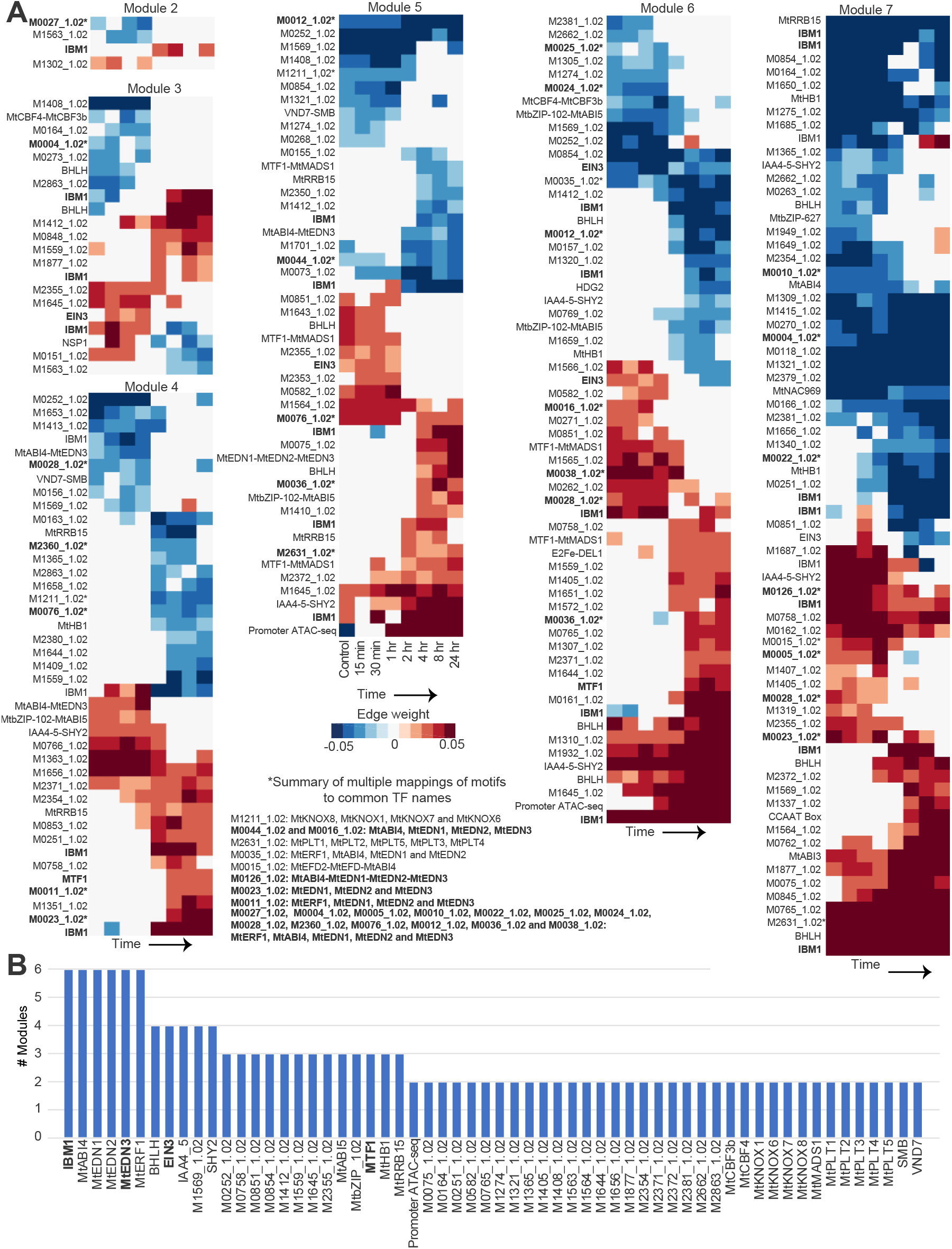
Regulator prioritization results. **(A)** Summary of DRMN regression weights that meet a T*-*test criterion of significant change (*P*<0.05) between 0-1 and 2-24 hr. CisBP motif IDs mapped to >=3 common names (*) are summarized separately (bottom center). **(B)** Regulators prioritized based on the frequency with which they are selected across modules with the T-test criteria. Labels of motifs mapped to IBM1, EDN3, MTF1, EIN3 are in bold in both panels.

### Identification of the targets of DRMN-prioritized regulators

DRMN identified regulators of gene expression dynamics in response to LCOs. Next, we aimed to identify their gene targets. Expression-based network inference is commonly used to define regulator-gene relationships (33), but is challenging with 8 time-points. To address this, we first obtained clusters of transitioning genes from DRMN modules (**Fig. 5D, Fig. S10A**), similar to those from ESCAROLE (**Fig. 2B, Fig. S10B**). We identified 79 transitioning gene clusters including 10,176 genes, of which 5,332 (>50%) were differentially expressed with DESeq (hypergeometric-test overlap *P*<0.05) and (8,398) 77% were identified in ESCAROLE as well, indicating consistency between the analyses. Next, we used two criteria based on (1) DRMN modules and motif enrichment, and (2) DRMN transitioning gene sets and regulatory motifs selected by a regularized regression method, Multi-Task Group LASSO (MTG-LASSO). The motif enrichment approach provides a more inclusive prediction of targets, while the regularized regression is more stringent. *Targets identified based on DRMN modules and motif enrichment*.

The first criterion to define a regulator’s targets was based on identifying genes belonging to a DRMN module predicted to be regulated by that regulator based on the DRMN regression models. Only genes containing the motif to which the regulator binds to in their promoter were considered targets. Moreover, only genes that belong to a module enriched for the associated motif were considered (*q*-value <0.05). This approach defined 3,888,784 edges connecting 298 regulatory motifs to 21,673 target genes and predicted targets of several DRMN-prioritized regulators, including IBM1, ERF1, EDN1-3, EIN3, SHY2, ABI4-5, MTF1 and KNOX and PLT family members.

#### Targets identified based on DRMN transitioning gene sets and MTG-LASSO

Our second approach modeled the variation in expression of each of the 79 transitioning gene clusters using a structured sparsity approach, Multi-Task Group LASSO (MTG-LASSO) (SLEP v4.1 package (34), **Fig. 7A**) to identify regulators (motifs/TFs) for each of the transitioning gene clusters. Here the same feature data from the DRMN analysis was used. We determined MTG-LASSO parameter settings for all 79 transitioning gene sets, identifying 33 with significant regulatory motif associations (**Fig. 7B, Fig. S11**). This generated 122,245 regulatory edges connecting 126 regulatory motifs to 5,978 genes (**Fig. 7C**).

**Figure 7.**
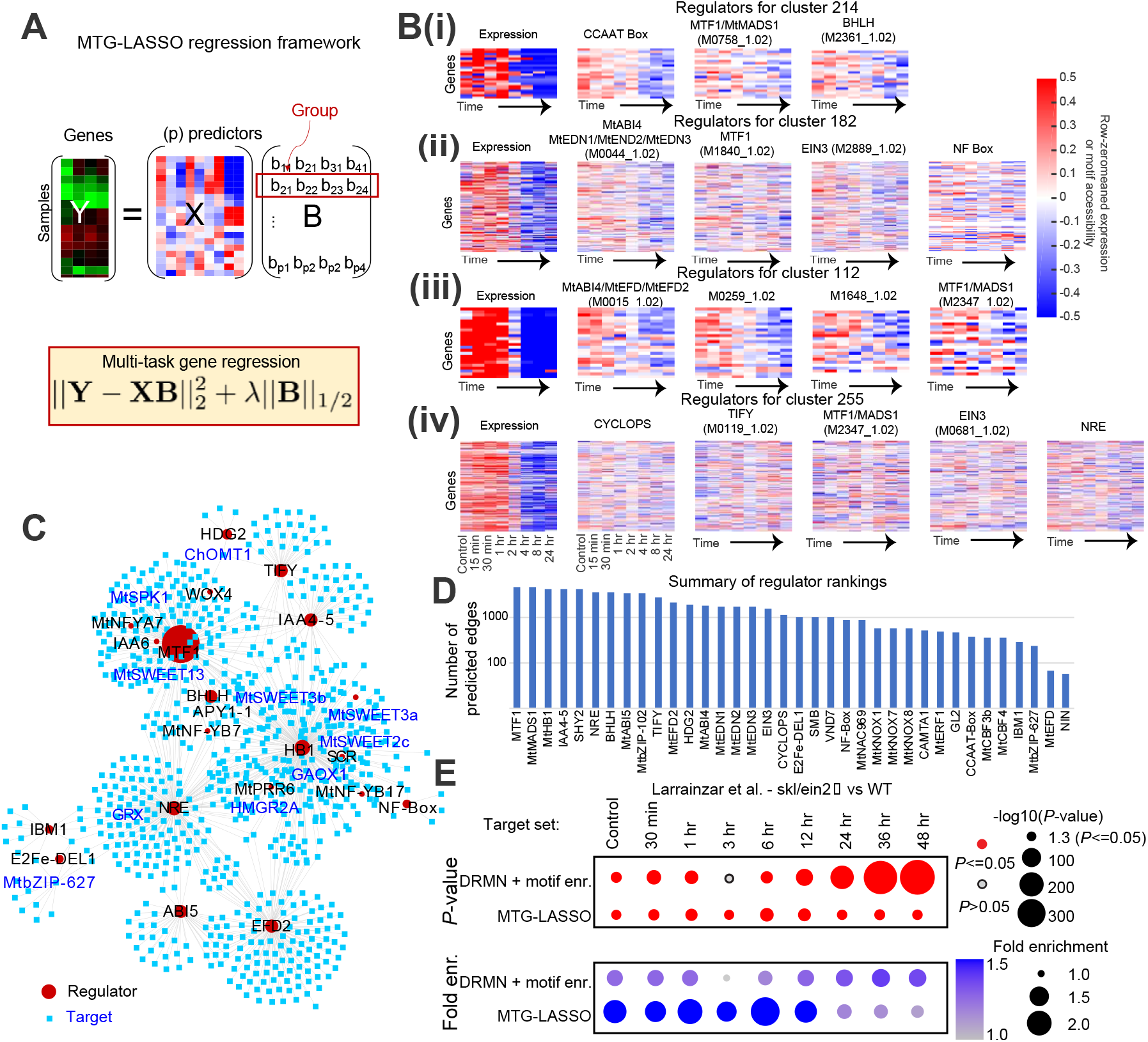
Multi-task group LASSO (MTG-LASSO) to predict regulators of transitioning genes. **(A)** MTG-LASSO was applied to infer significant regulatory features for each transitioning gene set. Shown is a model of predicting expression (**Y**) for a set of genes using the features (**X**) for the genes and coefficients (**B**). MTG-LASSO picks the same regulators for all genes in a set but allows for different regression weights. The regression weight of each regulator (row) is a group. **(B)** Example transitioning gene clusters presenting corresponding gene expression and motif accessibility profiles for regulators of interest (IBM1, MTF1, EIN3, EDN3). For each cluster, we show those genes presenting significant change in accessibility between 0-2 and 4-24 hr (T-test *P*-value <0.05) for at least one regulatory feature per cluster. **(C)** Visualization of the top 1000 predicted TF-gene network edges, ranked by regression weight magnitude from MTG-LASSO. **(D)** Ranking of regulators by the number of predicted targets highlights regulators identified at the module level (**Fig 6**), and additional ones like TIFY and CYCLOPS. **(E)** Overlap statistics of predicted EIN3 targets with DE genes between the *skl/ein2* and WT time-course data from Larrainzar et al.. Overlap is quantified at each time point with both hypergeometric test (-log10(*P*)) results and fold-enrichment scores (size and color).

Several gene sets exhibit consistent downregulation of expression and corresponding reduction in accessibility of predicted regulatory motifs between 0-2 and 4-24 hr (**Fig. 7B**). For example, gene set 214 (57 genes) shows downregulation of gene expression and reduced motif accessibility (after 4 hr) for multiple TFs: MTF1 and BHLH (**Fig. 7B, D**). Similarly, gene set 182 was predicted to be regulated by EDN3, MTF1, EIN3, and NF-Box motif, and exhibited correlated trends between gene expression and regulatory feature accessibility (**Fig. 7B**).

#### Computational validation of predicted targets by comparison with published data

To computationally validate our predicted targets, we compared our predicted EIN3 targets to differentially expressed (DE) genes from the Larrainzar et al. experiment comparing wild-type and *skl/ein2* mutant lines of *M. truncatula* (12). We selected *skl/ein2* because EIN2 and EIN3 are functionally related (35), and EIN3 was implicated as a significant regulator in the DRMN analysis. We find substantial overlap (hypergeometric test *P*<0.05) between the DE genes identified in mutant/wild-type comparisons at all time points, and our predicted targets (**Fig. 7E, Fig. S12A**). Moreover, when ranked by DESeq -log10(*P)* scores from the *skl/ein2* (relative to wild type) data, our predicted EIN3 targets are significantly (*P*<0.05) more differentially expressed than other genes (**Fig. S12B**) at all time points. This was observed using both Gene Set Enrichment Analysis (GSEA, (36, 37)) and Wilcoxon rank sum tests (**Supplementary Methods**). The significant overlap between DRMN/MTG-LASSO-predicted targets of EIN3 and the experimentally measured (DE) targets of EIN2 using multiple statistical approaches provides support for the predicted targets from DRMN.

### *EIN3* and *ERF1* are important regulators of root nodule symbiosis in *M. truncatula*

To experimentally test the involvement of DRMN prioritized transcription factors in root nodule symbiosis, we selected three TFs, *EIN3, ERF1* and *IAA4-5*. We knocked down the expression of the corresponding genes by RNAi and examined the nodulation phenotype in composite *M. truncatula* plants **(Methods)**. Knockdown of *MtrunA17Chr5g0440591* (*EIN3*) and *MtrunA17Chr1g0186741* (*ERF1*) significantly lowered the number of nodules produced on the RNAi roots **(Fig. 8A, S13A**, *P*<0.05 from an ANOVA test followed by Tukey’s HSD test post-hoc). Knockdown of *MtrunA17Chr1g0166011* (*IAA4-5*) did not alter nodulation relative to the empty vector (EV) control **(Fig. S13B)**. These nodules were all colonized by *S. meliloti* (**Fig. 8B**). Together, these results validate the role of *MtrunA17Chr5g0440591* (*EIN3*) and *MtrunA17Chr1g0186741* (*ERF1*) in rhizobium-legume symbiosis, as predicted by DRMN.

**Figure 8.**
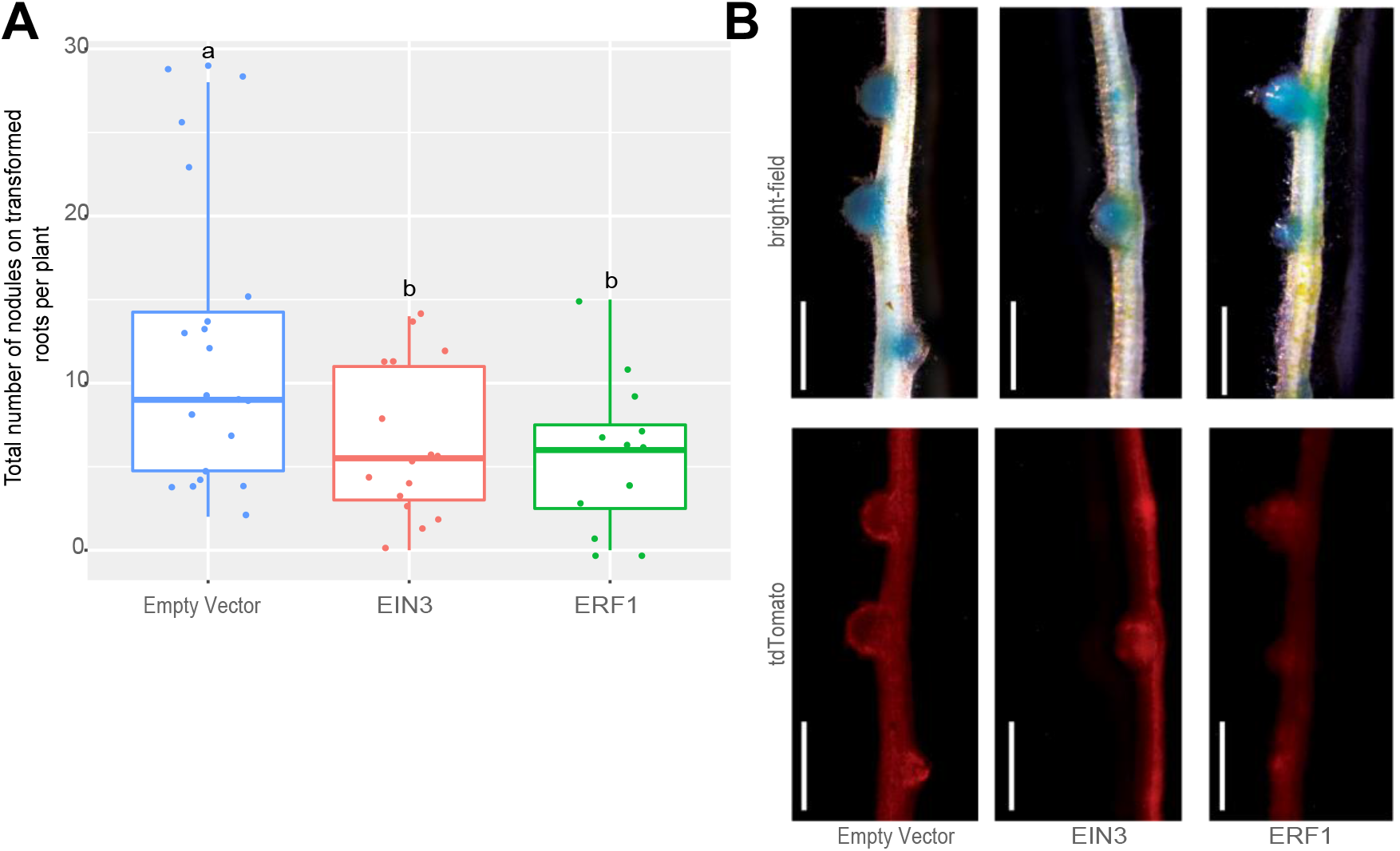
Knockdown of *MtrunA17Chr5g0440591* (*EIN3*) and *MtrunA17Chrg0186741* (*ERF1*) expression by RNA interference (RNAi) reduced the number of nodules on composite *M. truncatula* plants. **(A**) Data analyzed by ANOVA followed by Tukey’s HSD test for multiple comparisons. Box plots not connected by the same letter are significantly different (*P*<0.05). One extreme outlier (29 nodules) was excluded in the *MtrunA17Chrg0186741* (*ERF1*) experiment. (**B**) Images of nodules on subtending root supporting the effectiveness of RNAi. Blue color (top) indicates the rhizobial infection (*S. meliloti* constitutively expressing lacZ), and the red fluorescence marker (bottom) identifies transgenic roots (white scale bar = 1 mm).

## Discussion

The enormous economic and environmental cost of plant nitrogen fertilization motivates efforts towards identifying molecular mechanisms underlying legume perception of nitrogen-fixing bacteria and nodule development. Understanding such mechanisms is a pre-requisite for introducing this process in non-nitrogen-fixing species. We dissected the gene regulatory network in *M. truncatula* roots in response to *S. meliloti* LCOs by jointly profiling the temporal changes in the transcriptome and chromatin accessibility and integrating these data computationally. Extensive changes in the transcriptome are known to occur in *Medicago* roots in response to rhizobia signals and we show these changes are accompanied and facilitated by extensive chromatin remodeling. While the overall percentage of accessible chromatin regions remained similar across our time course experiment, regions of accessibility underwent a dramatic shift 1-2 hr after treatment. This remodeling appears to anticipate the development of root nodules, which requires stringent temporal and spatial control of gene expression. Chromatin accessibility of gene promoters notably also emerged as a significant predictor of gene expression (**Fig. 6**). These changes in chromatin accessibility enable and enhance the transcriptional changes required for nodule development by providing regulators access to promoters that may be inactive in other stages of plant development. Correlation was additionally observed between gene expression and promoter chromatin accessibility profiles of several essential regulators of nodulation, including *ERN1, CRE1, SKL/EIN2, IDP3/CYCLOPS*, and *ERN2*. Close coordination between chromatin accessibility and gene expression in LCO response is likely essential for root nodule development.

We applied novel methods for time-series analysis, ESCAROLE and DRMN (18), to model temporal changes in gene expression and chromatin accessibility. ESCAROLE enabled us to characterize the transcriptional dynamics beyond pairwise differential expression analysis, while DRMN allowed us to jointly analyze transcriptome and chromatin dynamics, and predict which transcription factors (TFs) are most important for expression dynamics. Consistent with the theme of chromatin reorganization under LCO treatment response, DRMN identified IBM1 as a critical regulator. IBM1 encodes a JmjC domain-containing histone demethylase that catalyzes the removal of H3K9 methylation and di-methylation in Arabidopsis (38). DRMN also identified regulatory genes involved in hormone responses in the early steps of symbiosis and nodule formation such as ethylene (*ERF1, EDN1-3* and *EIN3*) and ABA (*ABI4-5*). EIN3 is a transcription factor mediating ethylene-regulated gene expression and morphological responses in *Arabidopsis*. The role of EIN3 in rhizobium-legume symbiosis or LCOs signaling remains uncharacterized, but *sickle* (*skl*) mutants for an *EIN2* ortholog develop more infection threads and nodules and respond more to LCOs than wild-type plants, and ethylene treatment inhibits LCO signaling and nodule formation (39). ABI4 and ABI5, basic leucine zipper transcription factors implicated in several plant functions, coordinate LCO and cytokinin signaling during nodulation in *M. truncatula* (40). DRMN also identified regulators associated with the hormones involved in the nodule initiation, auxin (*SHY2*) and cytokinin (*MtRRB15*). *SHY2*, a member of the Aux/IAA family, plays a critical role in cell differentiation at root apical meristem and is activated by cytokinin (41, 42). *SHY2* was proposed as a candidate for nodule meristem regulation and differentiation after showing a very localized expression pattern in the nodule meristematic region (43). Also related to nodule meristem initiation, KNOX TF-family members and PLT1-5 were predicted as regulators of gene expression in response to LCOs. *MtPLT* genes (*MtPLT1-5*) are part of the root developmental program recruited from root formation, and control meristem formation and maintenance for root and nodule organogenesis (44). We experimentally validated two of our regulators, *EIN3* and *ERF1* using RNAi in *M. truncatula* and showed a significant effect in nodule formation. Prior work of Asamizu, *et al*. (45) independently supports the observation of the *ERF1* ortholog as an effector of nodule development in *L. japonicus*, where the number of nodules was likewise reduced in a similar RNAi experiment. Their findings suggest *ERF1* is induced by rhizobium on a 3 to 24 hr time scale, echoing the observed time scale of chromatin reorganization in *M. truncatula* in our work. Recent work of Reid, *et al*. (46) emphasizes an early, positive role of ethylene in rhizobium-legume symbiosis in *L. japonicus*, which supports why we observe ethylene-related TFs having a positive impact on nodulation, unlike the ethylene insensitive *skl* mutation (39). The large overlap observed between the potential EIN2 and EIN3 targets suggests they act through the same pathway during symbiosis. Based on this, we propose the early ethylene events promoting nodulation are mediated by the MtEIN3 copy identified in the present study, while late events (overridingly anti-nodulation) are mediated by EIN2. The exact mechanisms by which these genes regulate rhizobium-legume symbiosis can be explored in future research.

Finally, our analysis predicted genome-wide targets for transcription factors, including novel regulators identified by DRMN and previously known regulators of root nodulation, such as *NIN, NF-YA1/NF-YB1*, and *CYCLOPS*. This data generated several hypotheses about the regulation of gene expression in LCO response, informing future experimental validation of regulator-target relationships. For example, MTG-LASSO analysis predicted *NIN* as a direct target of SHY2 and MTF1, and *FLOT4*, required for infection thread formation, as a target of IBM1 (47). Among known regulators, MTG-LASSO indicated that *ARF16a* and *SPK1* are targets of NF-Y TFs. ARF16a and SPK1 control infection initiation and nodule formation (1). Several NF-Y genes (NF-YA5 and NF-YB17) were identified as regulated by CYCLOPS. In conclusion, our novel transcriptomic and accessibility datasets and computational framework to integrate these datasets provide a valuable resource to the plant community to understand the gene regulatory programs for the establishment of root nodulation symbiosis in *M. truncatula*.

## Materials and Methods

Detailed information, including plant material, growth conditions and treatments, ATAC-seq and RNA-seq library preparation and sequencing, differential expression analysis, ESCAROLE clustering, exploratory analysis of chromatin accessibility, Dynamic Regulatory Module Networks, and target prediction and validation analyses, are provided in Supplementary Information (SI) appendix (**Materials and Methods**). All data are described/linked in the SI Appendix.

The sequencing data presented in this publication have been deposited in NCBI’s Gene Expression Omnibus and are accessible through GEO Series accession number GSE154845 (https://www.ncbi.nlm.nih.gov/geo/query/acc.cgi?acc=GSE154845). The results from DRMN and MTG-LASSO gene target predictions are available in Zenodo (https://www.zenodo.org/deposit/4057115).

## Acknowledgments

This work was supported by the Department of Energy Office of Science Biological and Environmental Research (DE-SC0018247 to M.K. and J.-M.A.). We acknowledge the support of the Center of High Throughput Computing at UW-Madison to enable computational experiments.

