## Supplementary methods and figures for "Temporal change in chromatin accessibility predicts regulators of nodulation in *Medicago truncatula*"

**Supplementary Information Appendix:**

Materials and Methods

Figures S1 – S13

References

### MATERIALS AND METHODS

#### Plant material and treatment

Seeds of wild-type *Medicago truncatula* Jemalong A17 were sterilized and germinated in 1% agar plates, including 1µM GA3. Plates were stored at 4°C for three days in the dark and placed at room temperature overnight for germination. Seedlings were grown vertically for five days on a modified Fahraeus medium with no nitrogen (1), in a growth chamber (24°C, 16 h light/8h dark cycle, 70 µmol $m^{-2}$ $s^{-1}$ photosynthetic photon flux). LCOs were purified from *S. meliloti* strain 2011 as described previously (2). Next, seedling roots were immersed in a solution of purified LCOs ($10^{-8}$ M) or 0.005% ethanol solution (control) for 1 h. Roots were cut and immediately used for nuclei extraction and generation of ATAC-seq libraries (see below), or snap-frozen in liquid nitrogen for posterior RNA isolation and sequencing. Roots were collected at 0 h (control), 15 min, 30 min, 1 h, 2 h, 4 h, 8 h and 24 hr after LCOs treatment. Roots from seven plants were pooled for each of three biological replicates used in RNA sequencing, while roots from 15 plants were pooled for one replicate used in ATAC-seq, in each time point of the experiment.

#### ATAC-seq library preparation and sequencing

For ATAC-seq library preparation, we followed the protocol described previously (3) with modifications. Before nuclei isolation, all materials were precooled to 4°C. Briefly, roots were chopped for 2 min in 1 ml of pre-chilled lysis buffer (15 mM Tris-HCl pH7.5, 2mM EDTA, 20 mM NaCl, 80 mM KCl, 0.5 mM spermine, 15 mM 2-ME, 0.15 % TritonX-100) in a cold room. This step was repeated four times with a 1 min interval between repetitions. The homogenate was filtered through one layer of pre-wetted Miracloth, loaded on the surface of a 2 mL dense sucrose buffer (1.7 M sucrose, 10 mM Tris-HCl pH8.0, 2 mM $MgCl_2$, 5 mM 2-ME, 1 mM EDTA, 0.15 % Triton X100), and centrifuged (2400 g, 20 min at 4°C). The supernatant was removed, the nuclei were resuspended in 500 µl of lysis buffer and then filtered in 70 µm and 40 µm filters consecutively. The nuclei were then collected by centrifuging the solution at 1000g for 5 min at 4°C. After washing with 950 µl 1×TAPS buffer (10 mM TAPS-NaOH, pH8.0, 5 mM $MgCl_2$), the samples were centrifuged again at 1000 g for 5 min at 4°C. The supernatant was removed, leaving the nuclei suspended in approximately 10 µl of solution. Next, 1.5 µl of Tn5 transposase (Illumina FC-121-1030), 15 µl of Tagmentation buffer, and 13.5 µl of $ddH_20$ were added to the solution. The reaction was incubated at 37°C for 30 min. The product was purified using a QIAGEN MinElute PCR Purification kit and then amplified using Phusion DNA polymerase. 1 µl of the product was used in 10 µl qPCR cocktail with Sybr Green. Cycle number X was determined as the cycle were the ¼ of the maximum signal was reached. Then we amplified the rest of the product in a Phusion (NEB) PCR system with X-2 cycles (10 to 15 cycles, 50 µl of reaction). Amplified libraries were purified with AMPure beads (Beckman Coulter), and library concentrations were determined using a Qubit. Sequencing was carried out in an Illumina HiSeqX (2×150 cycles) at the HudsonAlpha Institute for Biotechnology (Huntsville, AL, USA).

**RNA-seq library preparation and sequencing**

For each RNA extraction, roots from 7 plants were pooled and ground while keeping the sample frozen. RNA extraction was performed as described previously (4). Libraries were prepared using 1 µg of RNA in the NEBNext® Ultra™ Directional RNA Library Prep Kit following the supplier's instructions (New England Biolabs, Ipswich, MA, USA). Sequencing was carried out with an Illumina HiSeq3000 (2×100 cycles) at the Interdisciplinary Center for Biotechnology Research at the University of Florida (Gainesville, FL, USA).

**RNA-seq data pre-processing and quality control**

Between 8.7-17.9 million, 2×100 bp reads were obtained after sequencing the 24 RNA-seq libraries. Reads were aligned with Kallisto (5) to the *M. truncatula* transcriptome (v5, (6), **Fig. S1**). The average of the alignment rates across time points was 87-95%. A total of 37,536 genes were detected with non-zero expression at any of the time points. The data were LOESS-normalized and processed with SLEUTH (7) for further analysis. Finally, TPM expression values were quantile-normalized and log-transformed before being used as input for further analysis. Principle components analysis was applied to these data in MATLAB (**Fig. 1A**). For comparative purposes, transcriptome time course data related to root nodulation (8) obtained from the *M. truncatul* wild-type reference accession Jemalong A17 and three mutants (*lyk3*, *nfp*, and *skl/ein2*), were analyzed using the same Kallisto/SLEUTH approach. The 144 samples characterized in that experiment presented alignment rates of 91-96%, except four outliers with rates of 73-88%. Analysis of this data set resulted in detecting 40,988 genes with non-zero expression, of which 36,298 were in common with the 37,536 identified in the present LCO-treatment experiment (**Fig S2**).

**Differential expression analysis of RNA-seq time course and comparison with existing data**

DESeq (9) was applied to both the data generated in the present work and previously published data sets for four rhizobial treatment (8). The expected count matrices of each data set were used as input to the DESeq algorithm, used in a default manner per the author recommendations. For each of the five time-course experiments, we assessed differential expression relative to control (time 0 h) for each later time point (**Fig. S2A**). An adjusted *P* threshold of 0.05 was applied to select differentially expressed (DE) genes for each time point in each experiment. The union of differentially expressed genes across time points was used for comparisons between datasets (**Fig. S2B**). For the Larrainzar et al. data set (8), we also identified differentially expressed genes between the three mutants (*lyk3*, *nfp*, and *skl/ein2*) relative to the wild-type reference (Jemalong A17) for matched time points (**Fig S2A**, right). As in the first analysis the union of genes identified at any time point defined the set of differentially expressed genes for the dataset. We quantified the degree of overlap between DE gene sets with an F-score, or harmonic mean, of the fraction of overlapping genes in each set using the union across all time points (**Fig. S2B)** as well as individual pairs of time points (**Fig. S2C**). For two sets of $N_1$ and $N_2$ genes, respectively, and $N_O$ in common between the two, the F-score is defined as:

$$F = 2\frac{\frac{N_O}{N_1}\frac{N_O}{N_2}}{\frac{N_O}{N_1}+\frac{N_O}{N_2}}$$

**Expression clustering analysis with ESCAROLE**

We analyzed the LCO-treatment time course data with ESCAROLE (10) to characterize the temporal changes in the transcriptome. We included 37,536 genes with at least one non-zero fragment count in at least one of the 24 experiments (3 replicates × 8 time points). Transcriptome data from each time point were grouped by k-means clustering and used as an input module assignment for the ESCAROLE algorithm (**Fig. 3**). The algorithm was run for 100 iterations with non-fixed covariance GMM clustering, and $k$=7 modules. The selection of $k$=7 was determined by the mean silhouette index per time point and overall BIC-corrected likelihood score (**Fig. S3A**, **B**). From ESCAROLE, we obtain a module assignment for each gene at each time point and identified sets of genes that transition in their module assignment across the eight time points (**Fig. 2B**).

We define transitioning gene sets from ESCAROLE results by grouping genes with a similar module transition profile agglomerative hierarchical clustering (**Fig. 3C, Fig. S3D**). The pairwise distance between genes used for this clustering approach was the fraction of mismatches in the module assignment across the (8-point) time course. The distance threshold (to determine the cut on the dendrogram for the hierarchical clustering) and the minimum number of genes in a cluster were the input parameters to define the transitioning gene clusters in this approach. In choosing settings for these parameters we tested different pairwise distance threshold values (those corresponding to 0-4 mismatches between module assignment profiles) and examined the resulting cluster sets for their size, overlap with differentially expressed genes, and enrichments of Gene Ontology (GO) and motif terms (**Fig. S10**). We chose a pairwise distance threshold of 0.26 in the hierarchical clustering analysis (corresponding to two mismatches across the 8-point time course) based on these results, and used those clusters with 10 or more genes to define the 112 transitioning gene sets from the ESCAROLE results.

**Exploratory analysis of ATAC-seq data**

*Data pre-processing.*

Each of the eight ATAC-seq libraries was paired-end sequenced twice, and 54 to 235 million reads were obtained from each sequencing library (**Fig. S4A** and **B**). The data were aligned with Bowtie 2 (11) to the *M. truncatula* v5 genome, with 46-75% of the data found mappable (alignable) to the reference genome. Properly paired fragments with a quality score of 3 or greater were then obtained with “samtools view -Sb -q3 -f2,” (Properly paired, **Fig. S4A**, **B**) and duplicate-removal was applied with “samtools rmdup” (12) to define the final library data sets utilized (Selected, **Fig S4A**, **B**).

Fragment length distributions of each time-point data set (**Fig. S4C**) present the expected ~10 bp DNA pitch but not nucleosome occupancy dependence first illustrated by Buenostro et al. (13). This is consistent with previously published plant ATAC-seq data from Blajic, et al. (14) (see Fig. 2A of Blajic et al.). The absence of nucleosome occupancy dependence can be in part due to aspects of the ATAC-seq protocol as implemented in plants versus mammals. Another explanation could be the large proportion of our reads mapping to promoter regions, which tend to be nucleosome depleted further explaining the diminished nucleosome pitch.

Peak calling was performed by applying MACS2 (15) to ATAC-seq data from each time point using the command:

macs2 callpeak -t <bam file> -n <Name> --format BAMPE -gsize=3.4e8

We mapped these peaks to genes if the center of a peak was within 10 kbp upstream and 1 kbp downstream of a gene transcription start site (TSS). Peaks called at each time point were merged across time points to generate a set of "universal peaks" using custom scripts (https://github.com/Roy-lab/PeakMergingCode; **Fig. S5A** and **B**) based on two criteria: 1) peaks from two different time points had a Jaccard score overlap of 0.9 or higher, and 2) the peak from one time point was contained within the peak detected in another time point. Annotations of the universal peak set (**Fig. 3D, Fig. S6C** and **G**) were generated in two steps: 1) annotating all peaks centered within 10 kbp upstream and 1 kbp downstream a gene TSS as "Promoter" peaks, and 2) using Homer annotatePeaks.pl tool (16) for all remaining peaks.

*Correlation and clustering analysis*

To enable quantitative comparison of the chromatin accessibility profiles across time, we aggregated the ATAC-seq read counts in two sets of genomic regions: (1) gene promoter regions (defined as 2 kbp upstream to 2 kbp downstream of a given gene TSS) and (2) universal peaks described above using custom scripts (https://github.com/Roy-lab/aggregateSignalRegion_nonLog). Briefly, we first generated the per-base pair coverage of fragments for each time point data set with Bedtools (17) using the command *bedtools genomecov <bam file> -bp -pc*. For each ±2 kbp gene promoter region, the average coverage per-bp was estimated, log-ratio transformed relative to the global genome-wide average per-bp coverage in the respective time point data set. The genome-wide average per-bp coverage was obtained by dividing the total coverage on any base pair by the length of the genome. The signals aggregated to the promoter were used for downstream principal components analysis (**Fig. 3C**, **Fig. S5A**). Similarly, the signal for the universal peaks was obtained by obtaining the mean per-bp coverage of the respective peak region and dividing that by the global average per-bp coverage. For both data sets, this was followed by quantile normalization across time points, providing a continuous measure of the accessibility of gene promoter and peak regions.

To evaluate the relationship between gene expression and either promoter or universal peak accessibility, we first performed a zero-mean transformation of each gene's expression profile and the corresponding accessibility profiles. Next, a Pearson correlation was estimated. To assess the association's significance, we generated a null distribution of correlations from 1,000 random permutations of the time points. We computed a *P*-value that estimates the probability of observing a correlation in the permuted data more significant in magnitude than an observed correlation. We treated positive and negative correlations separately. For the eight time points in this data set Pearson correlations were typically significant ($P$<=0.05) when > 0.50 or < -0.50. The zero-meaned promoter and universal peak accessibility profiles were clustered with k-means clustering, and the optimal settings for *k* were determined separately for each data set. In both cases, the silhouette index (computed with correlation distance metric) was used to select the optimal *k*. Here *k*=6 clusters were chosen for the promoter accessibility data (**Fig. 3A**, **Fig. S5B**). For the universal peak accessibility profile clusters, we additionally used enrichments for motifs within the clusters of peaks to determine the optimal setting of *k*=9 clusters (**Fig. S6D** and **E**). The clusters were enriched for motif instances of several known regulators (**Fig. S6F**). Furthermore, the peaks in clusters 1 and 2 were more likely to be annotated as intergenic regions than peaks in any other cluster (**Fig. S6G**).

**Integrative analysis of RNA-seq and ATAC-seq time course data using the Dynamic Regulatory Module Networks algorithm**

We applied a novel algorithm, Dynamic Regulatory Module Networks (DRMN, (18)), to our RNA-seq and ATAC-seq time course data set to identify *cis*-regulatory elements and transcription factors associated with genes that exhibit dynamic behavior. The inputs to this algorithm are the RNA-seq time series data, the number of expression modules and regulatory features for each time point derived from the ATAC-seq time course by examining the genomic region around a gene's TSS. The algorithm outputs gene expression modules (states) for each time point and their associated regulatory programs comprising the *cis*-regulatory elements that best predict gene expression of a particular module.

To obtain the *cis*-regulatory features for each gene, we used 333 *M. truncatula* motif position weight matrices from the CisBP v1.02 database (19) and seven curated motifs of interest (including those for NIN, CYCLOPS (CYC-RE), and NSP1 and other binding motifs). ATAC-seq activity was aggregated for those known motif instances in the manner described above for promoter and universal peak regions. Motif finding was done for each of the associated position weight matrices using the *pwmmatch.exact.r* script (from the PIQ pipeline (20)) using the default log-likelihood score threshold of 5. Motifs mapped to 10 kbp upstream to 1 kbp downstream of gene TSSs were assigned as potential features describing the corresponding gene's expression. For each gene, the accessibility of multiple instances of the same motif mapped to that gene was summed. Finally, the aggregated motif accessibility feature data were merged across the time course and quantile normalized. The normalized accessibility data for ±2 kbp promoter regions were also included as a predictive feature of gene expression.

The DRMN algorithm takes as input the number of modules, $k$ and uses either a regularized regression model, Fused Lasso (21), or a graph prior approach to learn regression models for each module, $k$, for all time points jointly. Here we used the Fused Lasso within DRMN, which has the following objective:

$$\min_{\Theta} \sum_{c} \left\| X_{c,k} - Y_{c,k}\Theta_{c,k}^{T} \right\|_2^2 + \rho_1 \|\Theta_k\|_1 + \sum_{c,c\prime} \rho_2 \left\| \Theta_{c,k} - \Theta_{c\prime,k} \right\|_1 + \rho_3 \|\Theta_k\|_{2,1}$$

Here $X_{c,k}$ is the $n_k\ X\ 1$ vector of expression levels for $n_k$ genes in modules $k$ for time point $c$, $Y_{c,k}\ is$ $n_k$ X $p$ motif-accessibility feature matrix corresponding to the same genes with regression coefficients $\Theta_{c,k}^{T}$, and which represent the quantified association of gene expression with individual regulatory motif features. Here $\Theta_k$ is the matrix of coefficients across time points. The sum over $c, c'$ represents the sum over pairs of consecutive time points. Specifically, here $\|.\|_1$ is the $l_1$ norm (sum of absolute values), $\|.\|_2$ is the $l_2$ norm (square-root of the sum of each value), and $\|.\|_{2,1}$ is the $l_{1,2}$ norm, i.e., the sum of the $l_2$ norm of the columns of the given matrix. Furthermore, $\rho_1$, $\rho_2$, and $\rho_3$ are hyper-parameters of the model that need to be tuned for optimal training and inference of DRMNs. These parameters represent 1) a sparsity penalty, 2) enforcing similarity of features for consecutive time points (*c*), and, 3) enforcing an overall similarity of feature selection across all time points. We used several criteria to determine these hyper-parameter settings. The most important is the Pearson correlation of actual and predicted expression in three-fold cross-validation settings to assess the resulting models' predictive power inferred for varied settings of the hyperparameters. Additionally, the quality of the clustering (silhouette index scores), the BIC-corrected likelihood score, and stability of predictive power in three-fold cross-validation (**Fig. S7A** and **C**) were considered. We first varied $\rho_1$ (values of 1, 5-60 in increments of 5, 75 and 100), and $\rho_2$ (values of 0-60 in increments of 5, and 75, and 100) independently and assessed the resulting predictive power for all models inferred. Predictive power generally monotonically decreased with increasing values of either parameter for values of $\rho_1$>10, while for $\rho_2$<=25 the clustering was often unstable, prone to switches. A choice was made for $\rho_1$=5 over $\rho_1$=1, since predictive power correlation was found to be marginally higher for $\rho_2$=30-60.

With the $\rho_1$ parameter fixed to 5, a second independent scan of $\rho_2$ and $\rho_3$ was performed, with 1) $\rho_2$ varied from 25-60, 75 and 100, and 2) $\rho_3$ scanned for values of 0-60 in increments of 5, 75 and 100. For settings of $\rho_3$=5-20 there tended to be unstable predictive power of the least expressed module, recovering comparable but not greater performance compared to results for $\rho_3$=0 or $\rho_3$>20, indicating no advantage for setting $\rho_3$>0. We considered the cross-validation predictive power, silhouette index of modules, similarity to ESCAROLE modules, in determining a setting for $\rho_2$ (**Fig. S7A**). Comparable performance was found for $\rho_2$=30-60, but $\rho_2$=45 and 50 maximized the mean three-fold cross-validation performance. We selected $\rho_2$=45, as it was the lower of the two settings to avoid unnecessarily high values for a hyper-parameter. Based on this assessment, the hyperparameter settings of $\rho_1$=5, $\rho_2$=45, and $\rho_3$=0 were taken for our final results.

We ran DRMN on our time-course data set for *k*=7 input modules, based on the optimal numbers of modules determined in the ESCAROLE analysis. Each module was predicted to have multiple regulators based on DRMN's fused regression model. To allow initial interpretation of the regulators, we filtered them as follows: 1) the magnitude of regulator-module edge-weights in at least one time point being greater than 0.02, and 2) the regulatory motif being enriched in the module (FDR corrected *q* value from hyper-geometric test, $q$<0.05) for all timepoints (**Fig. S8**). Interpretation of the modules was also made with GO enrichment analysis, using an FDR corrected hypergeometric test ($q$<0.05) to define significant enrichment (**Fig. S9**).

To identify module network edges that were significantly varying in time we first merged module network edge weights across time points per module and identified those edge weights that were significantly varying (t-test $P$<0.05 as implemented in MATLAB with the ttest2() function) across the 0-1 hr and 2-24 hr portions of the time course. The choice to compare across the 1->2 hr time point transition was motivated by the observation of module reorganization at this time window (**Fig. 5C**). Those regulatory edges found to be significantly varying are likely important at the module level of organization (**Fig. 6**).

To identify gene sets that transition in their expression due to changes in their predictive regulatory programs, we grouped genes that changed their DRMN inferred module assignment across time points using the same agglomerative hierarchical clustering approach applied in the ESCAROLE transitioning gene clustering analysis. We performed GO and motif enrichment on these gene sets as well to assess the optimal threshold for cutting the dendrogram (**Fig. S10A**). In total we identified 79 gene sets spanning 10,176 genes. These gene sets were further analyzed using a regularized regression approach, MTG-LASSO, to identify regulators for each gene set.

*Inferring coarse and fine-grained regulator-target interactions*

We used two approaches that leverage DRMN modules to predict regulator-target interactions: (i) target prediction based on DRMN modules and motif enrichment, (ii) target prediction analysis from DRMN transitioning gene sets with MTG-LASSO. The first and more inclusive criteria to define targets of a given regulator was based on identifying genes belonging to a module predicted to be regulated by that regulator using DRMN and containing a motif to which that regulator binds to in their promoter region. Moreover, only modules that were significantly enriched for the predicted regulator's associated motif were considered targets (Hypergeometric test $q$<0.05). This approach defined "coarse" module level gene regulatory networks.

Our second approach identified more fine-grained connections by predicting regulators for individual genes using transitioning gene sets and a structured sparsity approach called Multi-Task Group Lasso (MTG-LASSO, **Fig. 7A**). MTG-LASSO is a type of multi-task learning, where one performs a regression for multiple tasks simultaneously to share information among the tasks. Here each gene in the gene set is a task, and MTG-LASSO enables us to select the same regulator for all genes in the set but with different regression parameters. The regression weight defines the "group" in MTG-LASSO for the regulator for all genes in the set. MTG-LASSO selects or unselects entire groups of regression weights. The MTG-LASSO objective for each gene set is:

$$\min_{\Theta} \sum_{g} \frac{1}{2} \left\| X_g - \sum_{m} Y_{m,g}\, \Theta_{m,g} \right\|_2^2 + \lambda \|\Theta\|_{1/2}$$

Here $X_g$ is the expression profile over time for gene $g$, and $Y_{m,g}$ is the vector of motif accessibility features for motif $m$ and gene $g$ over time. The parameters $\Theta_{m,g}$ are the regression coefficients for predicting the expression of $g$ using the feature data for motif $m$. This second term denotes the $\|.\|_{1/2}$ norm defined as $\sum_m \sum_g \Theta_{m,g}^2$ and is used for 1) penalizing the number selected motif features according to the $l_1$ norm, and 2) enforcing smoothness of the regression coefficients across genes according to the $l_2$ norm. $\lambda$ is the hyper-parameter for controlling the group structure.

For each of the 79 transitioning gene sets, MTG-LASSO was applied (using the SLEP v4.1 package (22) in MATLAB (23)) to infer the most predictive regulatory features of gene expression over time from the same motif accessibility features used in the DRMN analysis. For each gene set we applied MTG-LASSO in a leave-one-out testing mode (**Fig. S11**), where each of the eight time points was left out one at a time, a model was fit on the remaining seven, and predictive power (Pearson's correlation) was computed on the left-out time point. For each regulator, we calculated a *P*-value to assess the significance of the frequency with which a given regulator was selected relative to random. This was achieved by randomizing the data 40 times and estimating a null distribution for the rate with which that regulator was selected across folds. A z-test *P*-value was then obtained for the result relative to random.

We called a regulator significant if it was selected at least 6 of 8 time-point folds, and the number of times it was selected was significantly higher (*t* test $P$ <0.05) relative to random for the frequency of selection across folds. MTG-LASSO's hyper-parameter, $\lambda$ was determined for each transitioning gene set from the range 0.20-0.99 (in intervals 0.10) based on 1) the mean Pearson correlation (predictive power) of the inferred regulatory features, 2) the number of regulators (5-15 for most gene sets) identified as significant such that the ratio of the said identified regulators to target genes being close to 0.05 (**Fig. S11**). This approach identified 33 gene sets (of the original 79) with predicted regulators. For the remaining transitioning gene sets, significant regulators tended not to be found based on our criteria. This could be either because the available predictive features were not good descriptions of the respective gene expression profiles or regulators are only obtained for only one or two settings of $\lambda$, hindering an appropriate assessment of results.

For each of the 33 gene sets for which we identified regulators using MTG-LASSO (**Fig. S11**), we created putative regulator-target predictions between the significant regulatory features and member genes, defining 122,245 regulatory edges spanning 126 regulatory features for 5,978 target genes (from 10,176 genes aggregated among the 79 transitioning gene clusters). Of the 126 motifs, we mapped 53 motifs to 278 *M. truncatula* regulator genes, including 31 well-studied regulators (specifically with common names in the v5 genome annotations). The remaining 73 motifs were assigned to 261 *M. truncatula* genes in the v5 genome assembly that were additionally identified as transcription factors (TFs). The relatively high number of motif to gene name mappings is because TF names were provided in CisBP v1.2 as systematic gene names from the v3/v3.5 *M. truncatula* genome assemblies rather than v5. We used a 70% BLAST similarity score to define mappings from *M. truncatula* v3/v3.5 genome systematic gene names to v5 genome systematic gene names.

*Comparison of DRMN targets and published regulator perturbation experiments*

To assess the validity of our predicted targets, we considered EIN3 and tested the overlap between two predicted target sets, (1) DRMN motif-enrichment-based and (2) MTG-LASSO-based targets, to differentially expressed gene sets from a previously published experiment comparing a mutant accession relative to wild-type plants (8). We mapped 6 CisBP motifs to EIN3 and used all of them in DRMN to predict targets. We obtained the differentially expressed gene sets from the *skl/ein2* mutant and WT time course data from Larraizar et al. (8). Briefly, we aligned the RNA-seq data to the *M. truncatula* v5 genome using Kallisto/SLEUTH and obtained differentially expressed genes at each time point using DESeq using an adjusted $P < 0.05$. We first used the Hypergeometric test with a threshold $P < 0.05$ to test for significant overlap for each of the predicted sets and differentially expressed genes for EIN2. Fold enrichment was calculated from the number of overlapping genes ($N_o$), the number of predicted targets from either the module level or MTG-LASSO per-gene level results ($N_T$) the number of identified differentially expressed genes ($N_{DE}$), and the "universe" of expressed (analyzed) genes ($N_U = 37{,}356$) as

$$Fold\ enrichment = \frac{N_O/N_T}{N_{DE}/N_U},$$

where the hypergeometric tests were applied correspondingly for the same numbers ($N_o, N_T, N_{DE}\ and\ N_U$) for each comparison. The comparisons from all timepoints present significant overlap (**Fig. 7E**, **Fig. S12A**), and indicate agreement between our predictions and the experimentally observed differentially expressed gene sets.

We further tested the degree to which predicted EIN3 targets were more differentially expressed relative to other genes in general in the Larrainzar et al. *skl/ein2* mutant condition (relative to WT) at each respective time point of that experiment. We ranked all genes by -log10(*P*) scores from DESeq applied to that data for each time point. The degree to which predicted EIN3 targets were more differentially expressed than other genes in that condition was assessed both with the Wilcoxon rank sum test (ranksum) function in MATLAB (23)) and the Gene Set Enrichment Analysis (GSEA) "Pre-ranked" algorithm (24, 25), again for each time-point. GSEA was applied in two ways: in a default weighted analysis with "meandiv" normalization of scores, and a "classic" mode with no normalization of scores, following the documentation and usage for this software. Both these tests show the predicted EIN3 targets are significantly differentially expressed than genes in general in the *skl/ein2* mutant condition, providing further support of the predicted target set of EIN3 (**Fig. S12B**,).

#### Validation of predicted regulators of nodulation with RNAi

We used RNAi to validate three predicted regulators from our DRMN analysis, EIN3, ERF1 and IAA4-5 104 bp region in the CDS specific to the gene of interest was amplified with 5'-CACC and inserted into pENTR™/D-TOPO® using directional TOPO® cloning, and further recombined *in-vitro* with the destination vector pK7GW1WG2(II)-RedRoot (https://gatewayvectors.vib.be/collection/pk7gwiwg2ii-redroot) using Gateway® LR Clonase® II enzyme mix using manufacturer's instructions.

To validate RNAi, total RNA was extracted from transformed roots of each genotype using Qiagen RNeasy® Plant Mini kit and genomic DNA removed using TURBO DNA-free™ Kit (Ambion). First-strand cDNA was synthesized using RevertAid RT Reverse Transcription Kit (Thermo Scientific™). Quantitative RT-PCR was performed using BIORAD SsoAdvanced Universal SYBR Green Supermix on BIORAD CFX96™ Real-time system; C1000 Touch™ Thermal cycler. The *HEL* and *UBC9* genes were used as endogenous controls. Two (*EIN3* - *MtrunA17Chr5g0440591*) or three (*ERF1* - *MtrunA17Chr1g0186741)* technical replicates were used. A BLAST was performed for all primers against the *Medicago v5* genome to ensure specificity. The primers chosen for the validation of RNAi do not overlap with the RNAi regions. All primers are listed in **Table S1**.

The RNAi expression clones were introduced into *Agrobacterium rhizogenes* MSU440 with electroporation. Composite *M. truncatula* plants were generated as previously described (26). Three weeks after transformation with *A. rhizogenes* MSU440, the roots were screened for red fluorescence of tdTomato, and the composite plants with red roots were transferred to growth pouches containing modified nodulation medium (MNM) (27). The plants were acclimated for 4 days and inoculated with *S. meliloti* 1021 harboring pXLGD4 (28). Two weeks post inoculation, live seedlings were stained for *lacZ* (5 mM potassium ferrocyanide, 5 mM potassium ferricyanide, and 0.08% X-gal in 0.1 M PIPES, pH 7) overnight at 37℃. Roots were rinsed in distilled water, and nodules were visualized and counted under a Leica fluorescence stereomicroscope.

**Table S1. Primers used in the RNAi validation study.**

| Purpose | Forward primer (5'-3') | Reverse primer (5'-3') |
|---|---|---|
| *MtrunA17Chr1g0166011* RNAi | **CACC**ATGGAATTCAAGGCAACTGAGCT | GTCTTTTGTTATTCTTAACAACACTACC |
| *MtrunA17Chr5g0440591* RNAi | **CACC**ATGATGATGTTTGAGGACATGGGG | CAGGCTCGGTTTGCCTGACAGAAGAAA |
| *MtrunA17Chr1g0186741* RNAi | **CACC**ATGGATTCAAGTTCAACCTCAA | CGTTCTCGTTGAAAGGAAGATAGTTG |
| Validation of *MtrunA17Chr5g0440591* RNAi | TAACGCAATCCCAGGAAAGAA | GCTGATAAGAGAGAACCCAAGG |
| Validation of *MtrunA17Chr1g0186741* RNAi | GAACGAACCAAGGAAGGAGAA | CCATTCCTCGTCGAATCTCTTATC |
| Endogenous control (*HEL)* | AGGACGCATGAGCTTTTCAA | AGCAGAGACCAGCATAACAAT |
| Endogenous control (*UBC9)* | AGCAGTGGTCTCATACTTGGA C | AGCCCCGCTTTGACAATATC |

SUPPLEMENTARY FIGURES

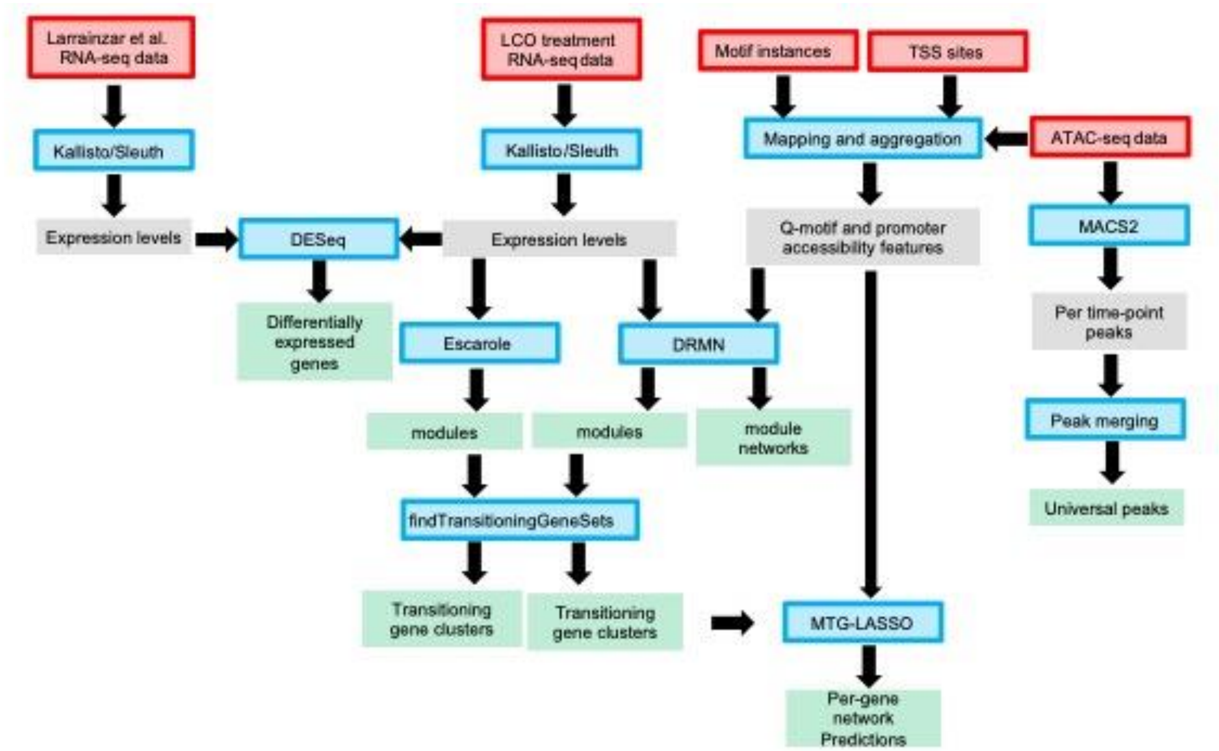


**Figure S1. Flow chart of the data analysis.** The input data are in red and the applied analysis algorithms and/or methods are in blue. Intermmediate and final analysis results are in gray and green, respectively.

**A**

Larrainzar et al. rhizobia treatment experiments

Present LCO treatment txperiment

| Time | Down | All | Up |
|---|---|---|---|
| 0.25 hr | 55 | 160 | 105 |
| 0.5 hr | 793 | 1,633 | 840 |
| 1 hr | 2,094 | 3,754 | 1,660 |
| 2 hr | 2,574 | 4,297 | 1,723 |
| 4 hr | 3,243 | 5,509 | 2,266 |
| 8 hr | 4,235 | 8,049 | 3,814 |
| 24 hr | 3,609 | 7,010 | 3,401 |
| Total | 7,052 | 12,839 | 7,540 |
| All pairs | 11,652 | 17,391 | 12,112 |

| | DE genes statistics (relative to control) | | | | | | | | | | | |
|---|---|---|---|---|---|---|---|---|---|---|---|---|
| | WT (A17) | | | *nfp* | | | *lyk3* | | | *skl* | | |
| Time | Down | All | Up | Down | All | Up | Down | All | Up | Down | All | Up |
| 0.5 hr | 186 | 1,033 | 847 | 18 | 200 | 182 | 30 | 420 | 390 | 15 | 289 | 274 |
| 1 hr | 399 | 1,247 | 848 | 11 | 107 | 96 | 34 | 316 | 282 | 57 | 410 | 353 |
| 3 hr | 489 | 1,045 | 556 | 168 | 365 | 197 | 267 | 563 | 296 | 226 | 497 | 271 |
| 6 hr | 764 | 1,595 | 831 | 311 | 551 | 240 | 485 | 1,031 | 546 | 609 | 1,373 | 764 |
| 12 hr | 536 | 1,372 | 836 | 321 | 720 | 399 | 414 | 938 | 524 | 967 | 2,185 | 1,218 |
| 24 hr | 166 | 751 | 585 | 0 | 7 | 7 | 29 | 352 | 323 | 2,314 | 5,245 | 2,931 |
| 36 hr | 632 | 1,525 | 893 | 261 | 532 | 271 | 448 | 1,025 | 577 | 3,888 | 7,326 | 3,438 |
| 48 hr | 185 | 786 | 601 | 11 | 126 | 115 | 34 | 375 | 341 | 4,650 | 8,895 | 4,245 |
| Total | 1,808 | 4,576 | 2,873 | 539 | 1,460 | 935 | 912 | 2,407 | 1,517 | 5,689 | 10,891 | 5,425 |
| All pairs | 6,090 | 8,987 | 6,315 | 1,852 | 2,925 | 1,973 | 3,015 | 4,513 | 2,988 | 8,984 | 15,252 | 7,925 |

| | DE gene statistics (relative to WT) | | | | | | | | |
|---|---|---|---|---|---|---|---|---|---|
| Time | Down | All | Up | Down | All | Up | Down | All | Up |
| t=0 | 23 | 33 | 10 | 10 | 25 | 15 | 67 | 142 | 75 |
| t=0.5 | 36 | 101 | 65 | 43 | 106 | 63 | 399 | 907 | 508 |
| t=1 | 110 | 248 | 138 | 143 | 203 | 60 | 296 | 715 | 419 |
| t=3 | 43 | 82 | 39 | 22 | 49 | 27 | 167 | 289 | 122 |
| t=6 | 91 | 474 | 383 | 17 | 47 | 30 | 334 | 725 | 391 |
| t=12 | 78 | 398 | 320 | 11 | 44 | 33 | 877 | 1,685 | 808 |
| t=24 | 89 | 311 | 222 | 26 | 82 | 56 | 1,834 | 3,455 | 1,621 |
| t=36 | 88 | 248 | 160 | 53 | 152 | 99 | 3,765 | 7,483 | 3,718 |
| t=48 | 182 | 370 | 188 | 65 | 201 | 136 | 3,959 | 8,017 | 4,058 |
| Total | 392 | 1,131 | 756 | 222 | 475 | 255 | 5,088 | 10,242 | 5,253 |

**B**

Comparison for down-regulated DE Genes

| | # DE genes | Present | WT | *nfp* | *lyk3* | *skl* | *nfp* vs. WT | *lyk3* vs. WT |
|---|---|---|---|---|---|---|---|---|
| Present | 7,052 | | | | | | | |
| WT | 1,808 | 0.22 | | | | | | |
| *nfp* | 539 | 0.11 | 0.37 | | | | | |
| *lyk3* | 912 | 0.15 | 0.52 | 0.55 | | | | |
| *skl* | 5,689 | 0.39 | 0.36 | 0.15 | 0.22 | | | |
| *nfp* vs. WT | 392 | 0.04 | 0.08 | 0.04 | 0.05 | 0.04 | | |
| *lyk3* vs. WT | 222 | 0.01 | 0.03 | 0.01 | 0.03 | 0.02 | 0.15 | |
| *skl* vs. WT | 5,088 | 0.13 | 0.08 | 0.03 | 0.04 | 0.05 | 0.05 | 0.03 |

F-score

Comparison for all DE genes

| # DE genes | Present | WT | *nfp* | *lyk3* | *skl* | *nfp* vs. WT | *lyk3* vs. WT |
|---|---|---|---|---|---|---|---|
| 12,839 | | | | | | | |
| 4,576 | 0.40 | | | | | | |
| 1,460 | 0.17 | 0.37 | | | | | |
| 2,407 | 0.26 | 0.56 | 0.51 | | | | |
| 10,891 | 0.53 | 0.43 | 0.20 | 0.29 | | | |
| 1,131 | 0.10 | 0.27 | 0.15 | 0.26 | 0.13 | | |
| 475 | 0.04 | 0.12 | 0.05 | 0.10 | 0.05 | 0.28 | |
| 10,242 | 0.48 | 0.37 | 0.15 | 0.22 | 0.72 | 0.15 | 0.06 |

F-score

Comparison for up-regulated DE genes

| # DE genes | Present | WT | *nfp* | *lyk3* | *skl* | *nfp* vs. WT | *lyk3* vs. WT |
|---|---|---|---|---|---|---|---|
| 7,540 | | | | | | | |
| 2,873 | 0.25 | | | | | | |
| 935 | 0.13 | 0.35 | | | | | |
| 1,517 | 0.18 | 0.57 | 0.48 | | | | |
| 5,425 | 0.33 | 0.39 | 0.20 | 0.29 | | | |
| 756 | 0.08 | 0.32 | 0.12 | 0.31 | 0.16 | | |
| 255 | 0.02 | 0.12 | 0.05 | 0.07 | 0.06 | 0.30 | |
| 5,253 | 0.27 | 0.24 | 0.10 | 0.15 | 0.08 | 0.10 | 0.03 |

F-score

**C**

Comparisons for down regulated DE genes (relative to control) — Per time point comparison with present data

| | Gene Stats | t=0->15 min | t=0->30 min | t=0->1 hr | t=0->2hr | t=0->4 hr | t=0->8hr | t=0->24 hr |
|---|---|---|---|---|---|---|---|---|
| | WT (A17) | | | | | | | |
| Gene stats | | 55 | 793 | 2,094 | 2,574 | 3,243 | 4,235 | 3,609 |
| t=0->30 min | 186 | 0.03 | 0.06 | 0.05 | 0.02 | 0.02 | 0.02 | 0.02 |
| t=0->1 hr | 399 | 0.02 | 0.11 | 0.09 | 0.04 | 0.03 | 0.03 | 0.03 |
| t=0->3 hr | 489 | 0.04 | 0.15 | 0.16 | 0.10 | 0.10 | 0.08 | 0.03 |
| t=0->6 hr | 764 | 0.02 | 0.16 | 0.18 | 0.12 | 0.13 | 0.13 | 0.06 |
| t=0->12 hr | 536 | 0.01 | 0.10 | 0.09 | 0.07 | 0.10 | 0.13 | 0.10 |
| t=0->24 hr | 166 | 0.01 | 0.02 | 0.02 | 0.02 | 0.02 | 0.01 | 0.02 |
| t=0->36 hr | 632 | 0.01 | 0.13 | 0.12 | 0.09 | 0.12 | 0.13 | 0.09 |
| t=0->48 hr | 185 | 0.01 | 0.03 | 0.03 | 0.02 | 0.02 | 0.01 | 0.02 |
| | *nfp* | | | | | | | |
| Gene stats | | 55 | 793 | 2,094 | 2,574 | 3,243 | 4,235 | 3,609 |
| t=0->30 min | 18 | 0.03 | 0.01 | 0.01 | 0.00 | 0.00 | 0.00 | 0.00 |
| t=0->1 hr | 11 | 0.03 | 0.00 | 0.00 | 0.00 | 0.00 | 0.00 | 0.00 |
| t=0->3 hr | 168 | 0.03 | 0.08 | 0.07 | 0.04 | 0.05 | 0.04 | 0.02 |
| t=0->6 hr | 311 | 0.03 | 0.11 | 0.11 | 0.06 | 0.09 | 0.09 | 0.04 |
| t=0->12 hr | 321 | 0.02 | 0.08 | 0.08 | 0.06 | 0.09 | 0.10 | 0.08 |
| t=0->24 hr | 0 | 0.00 | 0.00 | 0.00 | 0.00 | 0.00 | 0.00 | 0.00 |
| t=0->36 hr | 261 | 0.02 | 0.08 | 0.07 | 0.05 | 0.09 | 0.09 | 0.06 |
| t=0->48 hr | 11 | 0.00 | 0.00 | 0.00 | 0.00 | 0.00 | 0.00 | 0.00 |
| | *lyk3* | | | | | | | |
| Gene stats | | 55 | 793 | 2,094 | 2,574 | 3,243 | 4,235 | 3,609 |
| t=0->30 min | 30 | 0.05 | 0.01 | 0.01 | 0.00 | 0.00 | 0.00 | 0.00 |
| t=0->1 hr | 34 | 0.02 | 0.01 | 0.01 | 0.00 | 0.00 | 0.00 | 0.00 |
| t=0->3 hr | 267 | 0.03 | 0.12 | 0.11 | 0.06 | 0.07 | 0.06 | 0.02 |
| t=0->6 hr | 485 | 0.02 | 0.15 | 0.16 | 0.09 | 0.11 | 0.11 | 0.04 |
| t=0->12 hr | 414 | 0.01 | 0.11 | 0.10 | 0.06 | 0.10 | 0.12 | 0.08 |
| t=0->24 hr | 29 | 0.00 | 0.00 | 0.00 | 0.00 | 0.00 | 0.00 | 0.00 |
| t=0->36 hr | 448 | 0.02 | 0.12 | 0.10 | 0.07 | 0.11 | 0.12 | 0.08 |
| t=0->48 hr | 34 | 0.02 | 0.00 | 0.00 | 0.00 | 0.00 | 0.00 | 0.01 |
| | *skl* | | | | | | | |
| Gene stats | | 55 | 793 | 2,094 | 2,574 | 3,243 | 4,235 | 3,609 |
| t=0->30 min | 15 | 0.00 | 0.00 | 0.00 | 0.00 | 0.00 | 0.00 | 0.00 |
| t=0->1 hr | 57 | 0.00 | 0.03 | 0.02 | 0.01 | 0.01 | 0.01 | 0.00 |
| t=0->3 hr | 226 | 0.02 | 0.09 | 0.09 | 0.06 | 0.06 | 0.05 | 0.02 |
| t=0->6 hr | 609 | 0.02 | 0.15 | 0.17 | 0.11 | 0.13 | 0.12 | 0.06 |
| t=0->12 hr | 967 | 0.01 | 0.11 | 0.13 | 0.12 | 0.19 | 0.21 | 0.18 |
| t=0->24 hr | 2,314 | 0.01 | 0.11 | 0.16 | 0.16 | 0.19 | 0.20 | 0.22 |
| t=0->36 hr | 3,888 | 0.00 | 0.12 | 0.19 | 0.22 | 0.27 | 0.31 | 0.27 |
| t=0->48 hr | 4,650 | 0.01 | 0.11 | 0.19 | 0.21 | 0.23 | 0.25 | 0.24 |

Comparisons for all DE genes — Per time point comparison with present data

| | Gene Stats | t=0->15 min | t=0->30 min | t=0->1 hr | t=0->2hr | t=0->4 hr | t=0->8hr | t=0->24 hr |
|---|---|---|---|---|---|---|---|---|
| | WT (A17) | | | | | | | |
| Gene stats | | 160 | 1,633 | 3,754 | 4,297 | 5,509 | 8,049 | 7,010 |
| t=0->30 min | 1,033 | 0.05 | 0.22 | 0.18 | 0.15 | 0.18 | 0.14 | 0.14 |
| t=0->1 hr | 1,247 | 0.05 | 0.21 | 0.20 | 0.18 | 0.18 | 0.16 | 0.16 |
| t=0->3 hr | 1,045 | 0.04 | 0.15 | 0.17 | 0.16 | 0.16 | 0.13 | 0.13 |
| t=0->6 hr | 1,595 | 0.04 | 0.17 | 0.20 | 0.21 | 0.22 | 0.18 | 0.17 |
| t=0->12 hr | 1,372 | 0.04 | 0.15 | 0.16 | 0.16 | 0.17 | 0.16 | 0.16 |
| t=0->24 hr | 751 | 0.04 | 0.08 | 0.08 | 0.08 | 0.09 | 0.07 | 0.08 |
| t=0->36 hr | 1,525 | 0.03 | 0.14 | 0.16 | 0.16 | 0.17 | 0.15 | 0.16 |
| t=0->48 hr | 786 | 0.03 | 0.06 | 0.07 | 0.08 | 0.08 | 0.06 | 0.08 |
| | *nfp* | | | | | | | |
| Gene stats | | 160 | 1,633 | 3,754 | 4,297 | 5,509 | 8,049 | 7,010 |
| t=0->30 min | 200 | 0.03 | 0.07 | 0.04 | 0.04 | 0.04 | 0.03 | 0.03 |
| t=0->1 hr | 107 | 0.06 | 0.04 | 0.02 | 0.03 | 0.03 | 0.02 | 0.02 |
| t=0->3 hr | 365 | 0.04 | 0.08 | 0.08 | 0.07 | 0.07 | 0.06 | 0.05 |
| t=0->6 hr | 551 | 0.03 | 0.09 | 0.10 | 0.09 | 0.11 | 0.09 | 0.07 |
| t=0->12 hr | 720 | 0.04 | 0.09 | 0.11 | 0.10 | 0.10 | 0.10 | 0.11 |
| t=0->24 hr | 7 | 0.01 | 0.00 | 0.00 | 0.00 | 0.00 | 0.00 | 0.00 |
| t=0->36 hr | 532 | 0.03 | 0.08 | 0.08 | 0.07 | 0.08 | 0.08 | 0.08 |
| t=0->48 hr | 126 | 0.03 | 0.02 | 0.01 | 0.02 | 0.02 | 0.01 | 0.02 |
| | *lyk3* | | | | | | | |
| Gene stats | | 160 | 1,633 | 3,754 | 4,297 | 5,509 | 8,049 | 7,010 |
| t=0->30 min | 420 | 0.07 | 0.14 | 0.09 | 0.07 | 0.08 | 0.07 | 0.07 |
| t=0->1 hr | 316 | 0.06 | 0.09 | 0.07 | 0.06 | 0.07 | 0.06 | 0.06 |
| t=0->3 hr | 563 | 0.05 | 0.11 | 0.11 | 0.10 | 0.10 | 0.08 | 0.08 |
| t=0->6 hr | 1,031 | 0.04 | 0.15 | 0.16 | 0.16 | 0.16 | 0.13 | 0.13 |
| t=0->12 hr | 938 | 0.04 | 0.13 | 0.14 | 0.12 | 0.14 | 0.13 | 0.12 |
| t=0->24 hr | 352 | 0.05 | 0.05 | 0.04 | 0.05 | 0.05 | 0.03 | 0.04 |
| t=0->36 hr | 1,025 | 0.04 | 0.12 | 0.13 | 0.13 | 0.14 | 0.12 | 0.12 |
| t=0->48 hr | 375 | 0.05 | 0.05 | 0.04 | 0.05 | 0.05 | 0.04 | 0.05 |
| | *skl* | | | | | | | |
| Gene stats | | 160 | 1,633 | 3,754 | 4,297 | 5,509 | 8,049 | 7,010 |
| t=0->30 min | 289 | 0.09 | 0.12 | 0.08 | 0.06 | 0.06 | 0.05 | 0.05 |
| t=0->1 hr | 410 | 0.07 | 0.12 | 0.10 | 0.09 | 0.09 | 0.07 | 0.07 |
| t=0->3 hr | 497 | 0.05 | 0.11 | 0.11 | 0.09 | 0.10 | 0.07 | 0.07 |
| t=0->6 hr | 1,373 | 0.04 | 0.15 | 0.18 | 0.18 | 0.20 | 0.16 | 0.15 |
| t=0->12 hr | 2,185 | 0.04 | 0.14 | 0.19 | 0.19 | 0.22 | 0.22 | 0.21 |
| t=0->24 hr | 5,245 | 0.02 | 0.12 | 0.20 | 0.22 | 0.24 | 0.28 | 0.29 |
| t=0->36 hr | 7,326 | 0.02 | 0.15 | 0.25 | 0.27 | 0.31 | 0.36 | 0.35 |
| t=0->48 hr | 8,895 | 0.01 | 0.13 | 0.24 | 0.27 | 0.29 | 0.35 | 0.35 |

Comparisons for up regulated DE genes (relative to control) — Per time point comparison with present data

| | Gene Stats | t=0->15 min | t=0->30 min | t=0->1 hr | t=0->2hr | t=0->4 hr | t=0->8hr | t=0->24 hr |
|---|---|---|---|---|---|---|---|---|
| | WT (A17) | | | | | | | |
| Gene stats | | 105 | 840 | 1,660 | 1,723 | 2,266 | 3,814 | 3,401 |
| t=0->30 min | 847 | 0.04 | 0.21 | 0.17 | 0.04 | 0.02 | 0.02 | 0.03 |
| t=0->1 hr | 848 | 0.04 | 0.18 | 0.16 | 0.05 | 0.03 | 0.02 | 0.03 |
| t=0->3 hr | 556 | 0.04 | 0.10 | 0.12 | 0.15 | 0.15 | 0.09 | 0.06 |
| t=0->6 hr | 831 | 0.05 | 0.13 | 0.15 | 0.17 | 0.20 | 0.14 | 0.08 |
| t=0->12 hr | 836 | 0.04 | 0.12 | 0.11 | 0.13 | 0.16 | 0.14 | 0.12 |
| t=0->24 hr | 585 | 0.05 | 0.09 | 0.09 | 0.10 | 0.11 | 0.08 | 0.09 |
| t=0->36 hr | 893 | 0.03 | 0.08 | 0.08 | 0.11 | 0.13 | 0.11 | 0.12 |
| t=0->48 hr | 601 | 0.04 | 0.06 | 0.07 | 0.08 | 0.08 | 0.06 | 0.09 |
| | *nfp* | | | | | | | |
| Gene stats | | 105 | 840 | 1,660 | 1,723 | 2,266 | 3,814 | 3,401 |
| t=0->30 min | 182 | 0.03 | 0.08 | 0.04 | 0.01 | 0.00 | 0.00 | 0.01 |
| t=0->1 hr | 96 | 0.06 | 0.03 | 0.02 | 0.01 | 0.00 | 0.00 | 0.00 |
| t=0->3 hr | 197 | 0.03 | 0.05 | 0.06 | 0.07 | 0.08 | 0.05 | 0.03 |
| t=0->6 hr | 240 | 0.01 | 0.03 | 0.04 | 0.06 | 0.10 | 0.08 | 0.04 |
| t=0->12 hr | 399 | 0.00 | 0.01 | 0.02 | 0.03 | 0.07 | 0.08 | 0.11 |
| t=0->24 hr | 7 | 0.00 | 0.00 | 0.00 | 0.00 | 0.00 | 0.00 | 0.00 |
| t=0->36 hr | 271 | 0.02 | 0.02 | 0.02 | 0.03 | 0.06 | 0.06 | 0.07 |
| t=0->48 hr | 115 | 0.03 | 0.03 | 0.02 | 0.03 | 0.04 | 0.02 | 0.03 |
| | *lyk3* | | | | | | | |
| Gene stats | | 105 | 840 | 1,660 | 1,723 | 2,266 | 3,814 | 3,401 |
| t=0->30 min | 390 | 0.06 | 0.17 | 0.12 | 0.02 | 0.01 | 0.01 | 0.02 |
| t=0->1 hr | 282 | 0.06 | 0.10 | 0.08 | 0.02 | 0.01 | 0.01 | 0.01 |
| t=0->3 hr | 296 | 0.05 | 0.08 | 0.10 | 0.10 | 0.10 | 0.05 | 0.03 |
| t=0->6 hr | 546 | 0.04 | 0.10 | 0.11 | 0.14 | 0.16 | 0.11 | 0.07 |
| t=0->12 hr | 524 | 0.04 | 0.09 | 0.09 | 0.11 | 0.15 | 0.12 | 0.11 |
| t=0->24 hr | 323 | 0.05 | 0.07 | 0.06 | 0.07 | 0.09 | 0.05 | 0.06 |
| t=0->36 hr | 577 | 0.04 | 0.07 | 0.07 | 0.08 | 0.12 | 0.09 | 0.11 |
| t=0->48 hr | 341 | 0.05 | 0.06 | 0.06 | 0.06 | 0.08 | 0.05 | 0.08 |
| | *skl* | | | | | | | |
| Gene stats | | 105 | 840 | 1,660 | 1,723 | 2,266 | 3,814 | 3,401 |
| t=0->30 min | 274 | 0.05 | 0.13 | 0.08 | 0.01 | 0.01 | 0.00 | 0.01 |
| t=0->1 hr | 353 | 0.03 | 0.09 | 0.07 | 0.02 | 0.02 | 0.01 | 0.02 |
| t=0->3 hr | 271 | 0.06 | 0.09 | 0.10 | 0.10 | 0.10 | 0.05 | 0.03 |
| t=0->6 hr | 764 | 0.04 | 0.11 | 0.11 | 0.18 | 0.23 | 0.16 | 0.11 |
| t=0->12 hr | 1,218 | 0.03 | 0.08 | 0.09 | 0.11 | 0.18 | 0.17 | 0.18 |
| t=0->24 hr | 2,931 | 0.01 | 0.06 | 0.08 | 0.08 | 0.14 | 0.20 | 0.22 |
| t=0->36 hr | 3,438 | 0.01 | 0.06 | 0.09 | 0.09 | 0.18 | 0.26 | 0.24 |
| t=0->48 hr | 4,245 | 0.01 | 0.05 | 0.09 | 0.08 | 0.15 | 0.22 | 0.22 |

**Figure S2. Summary of differentially expressed (DE) gene statistics. (A)** Summary statistics (red) of DE genes identified with DESeq (adjusted-*P*<0.05) in the LCO-treatment time course data set and the four rhizobial-treatment Larrainzar et al. time course data sets (WT, *nfp*, *lyk3*, and *skl*). Statstics are shown for all DE genes, and for specifically up- or down-regulated genes (relative to control or WT conditions). The Larrainzar et al. data results are shown for all time points both relative to control (within all four experiments) and relative to WT (for the three mutant strains). The overlap (F-score, blue color scale) of the identified gene sets **(B)** across time-points, and **(C)** per-time-point are also shown (see **Methods**). The *skl/ein2* mutant data DE gene set from Larrainzar et al. is the most similar to that of our present experiment, suggesting LCO-response is predominant in both experiments.

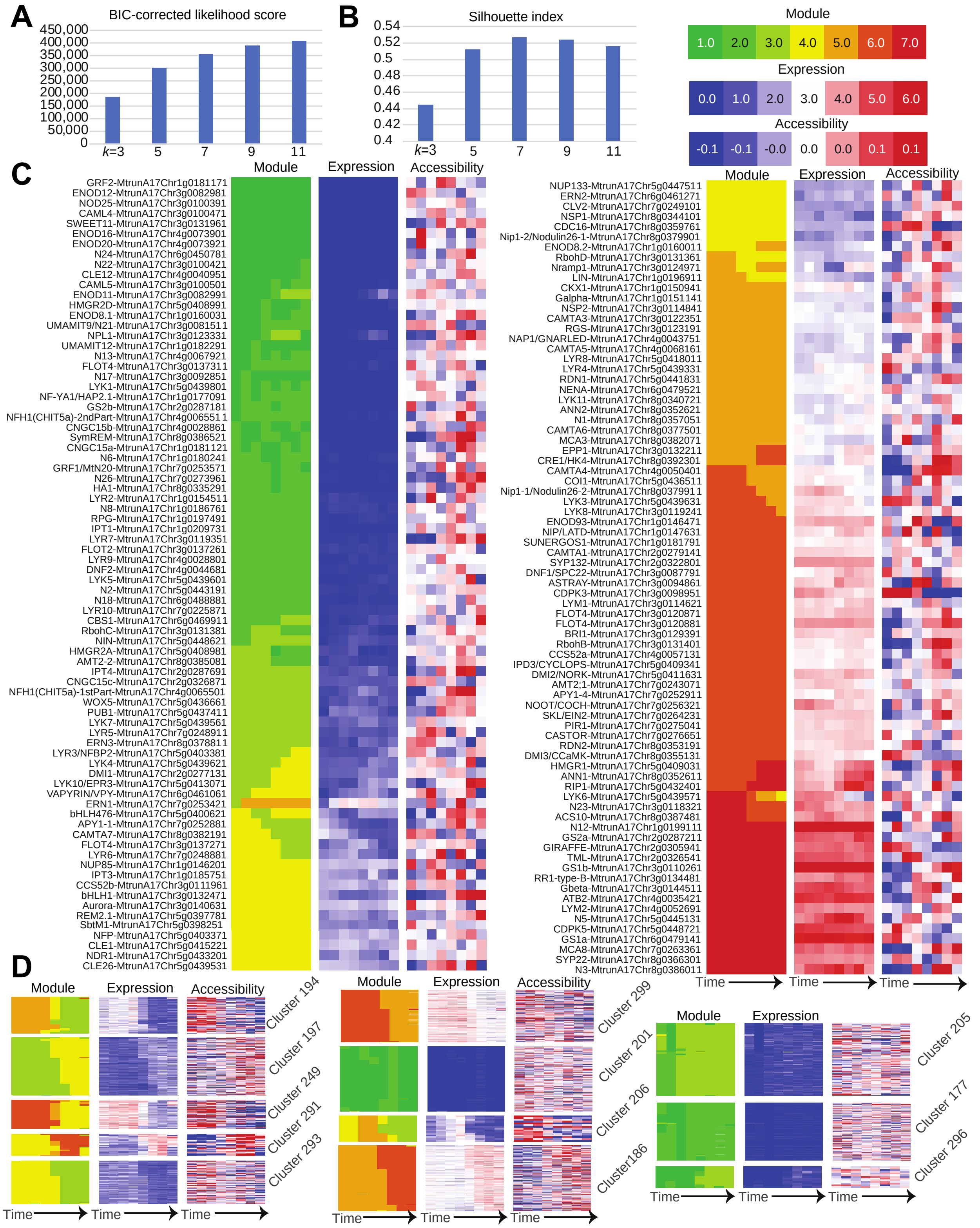


**Figure S3. ESCAROLE analysis of RNAseq data. (A)** Silhouette index, **(B)** and BIC-corrected likelihood scores for ESCAROLE results for varying settings of *k*, motivating *k*=7 as an appropriate choice for our analysis. **(C)** Summary of ESCAROLE module assignments, expression levels and row-zero-meaned promoter accessibility profiles for 152 genes of interest in root nodulation and

N-fixing symbiosis (see legend, upper right-hand corner). **(D)** Example transitioning gene sets identified with ESCAROLE module assignments. Shown are gene expression profiles, as well as promoter accessibility profiles (see legend, upper right-hand corner). Although not utilized in the inference of these clusters, promoter accessibility presents a clear association with gene expression for these gene sets.

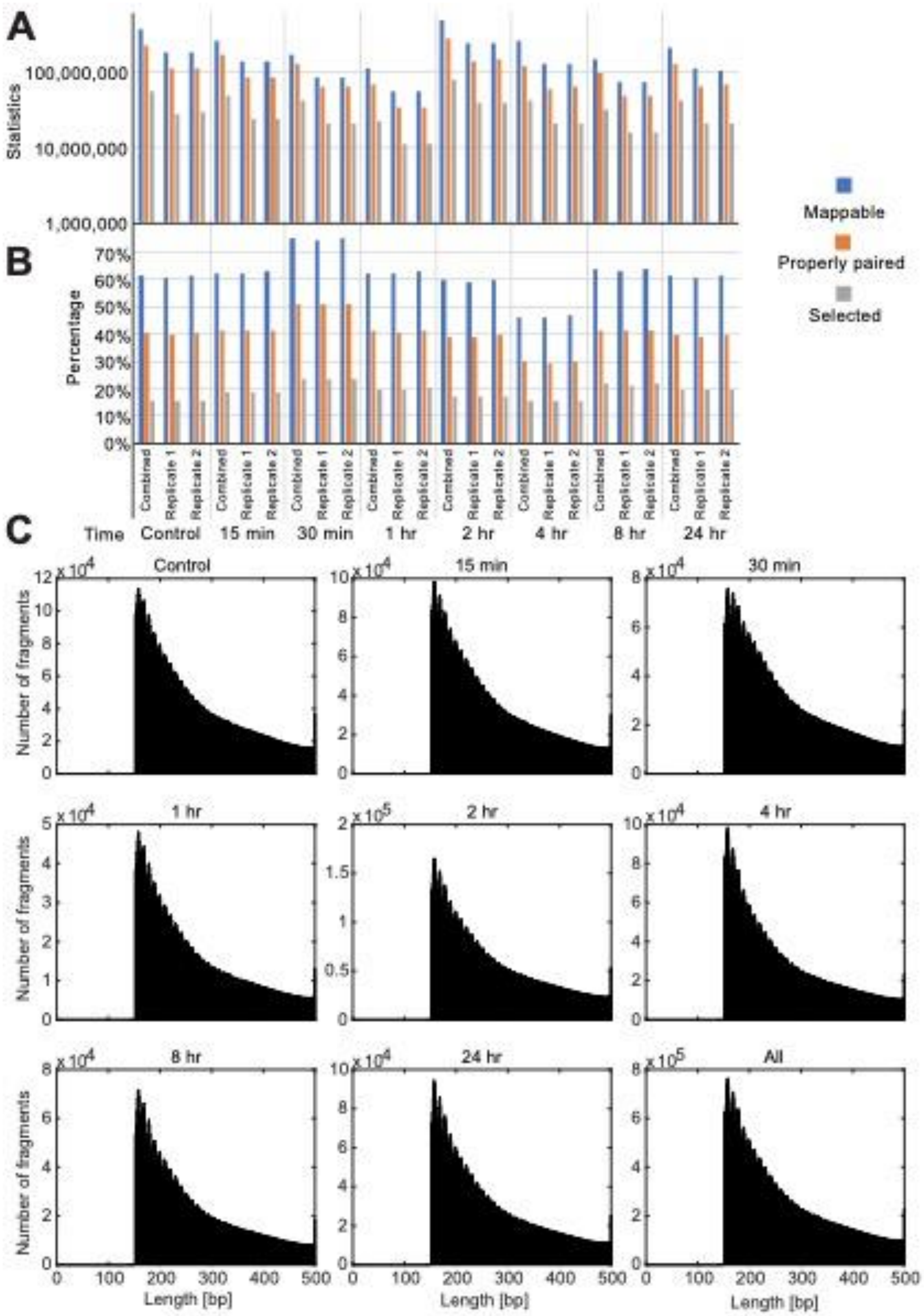


**Figure S4. Summary of ATACseq data alignment. (A)** Alignment statistics and **(B)** percentages of the data that are 1) mappable to the *M. truncatula* genome, 2) properly paired and 3) selected across replicates and time-points. The “selected” alignments are those meeting a mapping quality condition, and duplicate removal as described in the **Methods**. **(C)** Paired-end fragment length distributions shown for each time-point data set (and overall). Although nucleosome occupancy dependence is not prominent in these distributions, due to the specifics of the protocol implemented in this (plant) experiment, the ~10 bp DNA pitch is present.

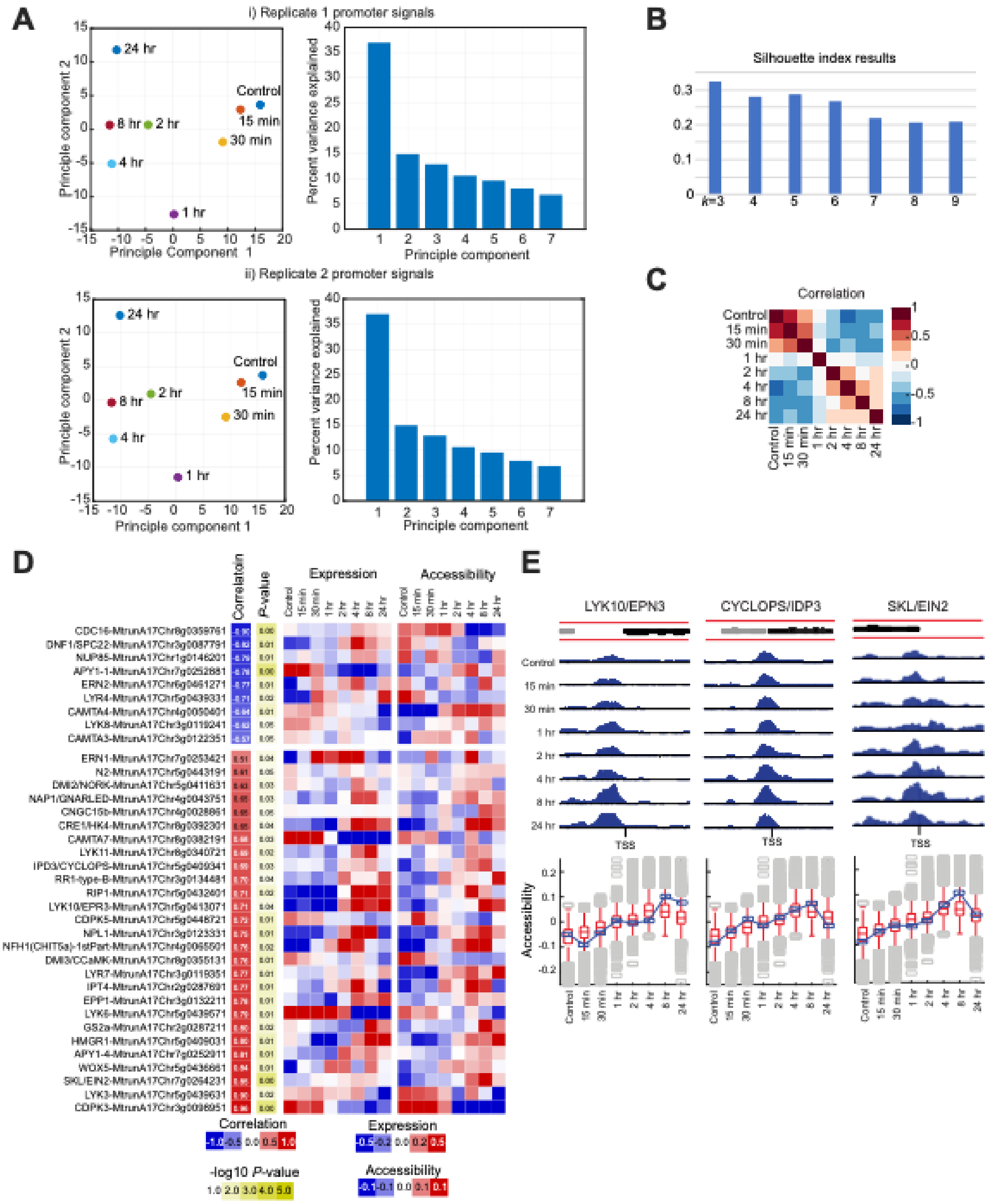


**Figure S5. Gene promoter chromatin accessibility (ATAC-seq) analysis. (A)** Principal Components Analysis (PCA) results of aggregated (±2 kbp of TSS) promoter signals for the ATAC-seq replicate data sets analyzed separately. PCA shows similar trends as in the combined data (**Fig. 3D**). **(B)** Silhouette index (SI) for k-means clustering applied to the row-zero-meaned promoter accessibility data, motivating a choice of *k*=6 clusters after which SI sharply decreases.

**(C)** Pearson correlation of the promoter accessibility data (with row-zero-mean transformation) across time points. **(D)** Row-zero-meaned gene expression and promoter accessibility profiles of 36 nodulation genes of interest with significant positive or negative correlation. **(E)** Normalized coverage tracks for promoter regions (±2 kbp) for known regulatory genes of interest (prepared in part with IGV (29)). In black are shown the genomic coordinates of the genes of interest, and in gray are nearby genes (if present).

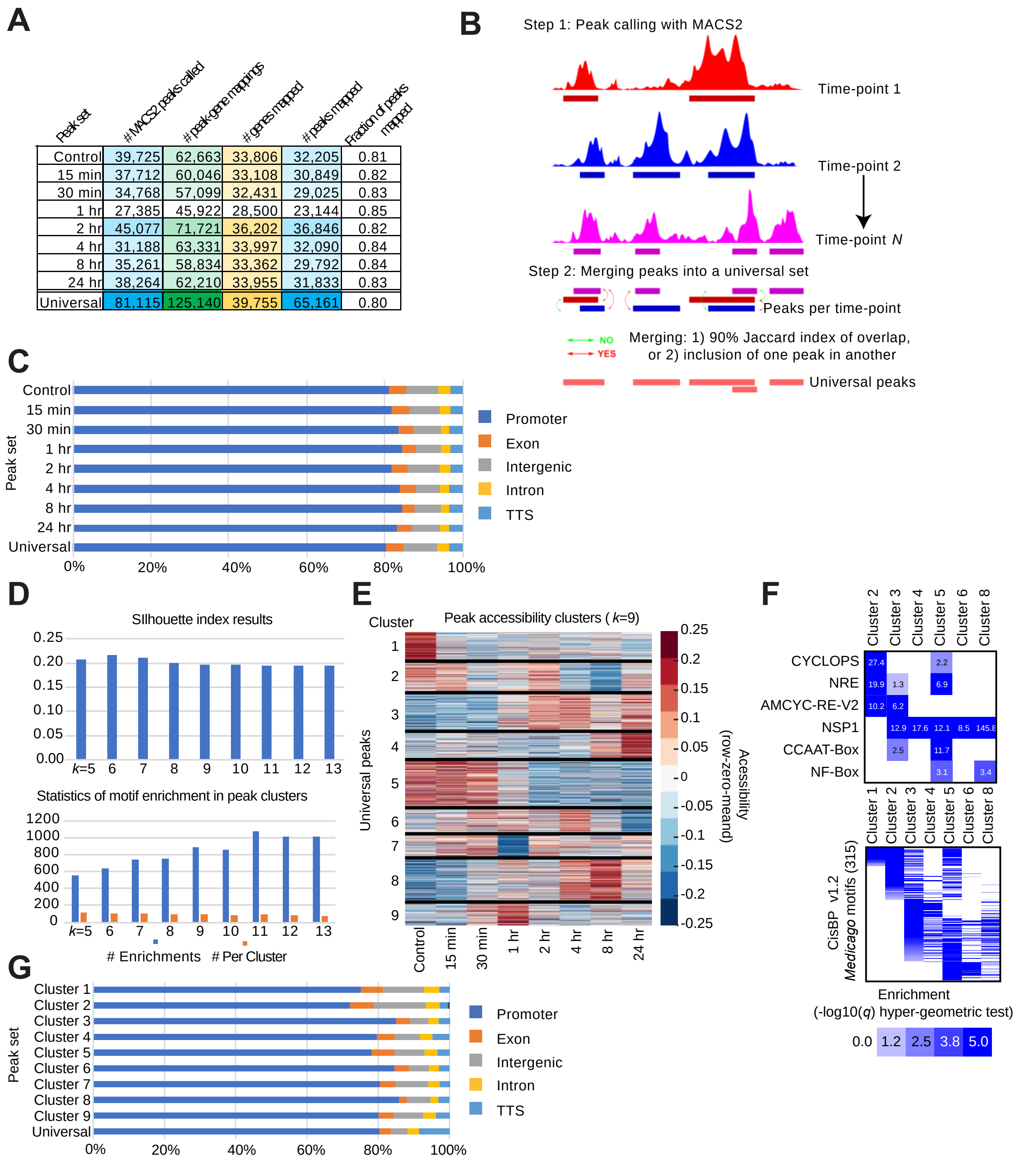


| Peak set | # MACS2 peaks called | # peak-gene mappings | # genes mapped | # peaks mapped | Fraction of peaks mapped |
|---|---|---|---|---|---|
| Control | 39,725 | 62,663 | 33,806 | 32,205 | 0.81 |
| 15 min | 37,712 | 60,046 | 33,108 | 30,849 | 0.82 |
| 30 min | 34,768 | 57,099 | 32,431 | 29,025 | 0.83 |
| 1 hr | 27,385 | 45,922 | 28,500 | 23,144 | 0.85 |
| 2 hr | 45,077 | 71,721 | 36,202 | 36,846 | 0.82 |
| 4 hr | 31,188 | 63,331 | 33,997 | 32,090 | 0.84 |
| 8 hr | 35,261 | 58,834 | 33,362 | 29,792 | 0.84 |
| 24 hr | 38,264 | 62,210 | 33,955 | 31,833 | 0.83 |
| Universal | 81,115 | 125,140 | 39,755 | 65,161 | 0.80 |

**Figure S6. Summary of ATAC-seq peak calling and accessibility analysis. (A)** MACS2 peak calling statistics. **(B)** Peak merging procedure to define a universal peak set across time points. Peaks are merged using the criteria of 90% Jaccard distance overlap or one peak being contained in another. **(C)** Summary of genomic annotations of peaks per time point and for the universal set. **(D)** Silhouette index scores and numbers of motif enrichments for k-means clusters of accessibility profiles aggregated for the universal peaks. **(E)** Temporal patterns of change in accessibility for the universal peak regions for *k*=9 clusters. **(F)** Enrichment (-log(*q*-value)) for motifs of interest (top) and general CisBP motifs (bottom) for the universal peak accessibility clusters connect putative regulators to the temporal patterns of change in accessibility. **(G)** Genomic annotations of peaks by cluster assignment. Clusters 1 and 2 show a higher proportion of peaks annotated to intergenic regions (11.6 and 14.8 %, respectively) compared to other clusters (5.3-9.2%), indicative of distal regulation.

**A**

| | Correlation | | | | Comparison to ESCAROLE modules | | | | | | | | | Silhouette index of modules | | | | | | | | |
|---|---|---|---|---|---|---|---|---|---|---|---|---|---|---|---|---|---|---|---|---|---|---|
| Setting (p1=5, and p3=0) | Full data set | CV training | CV test - mean across folds | CV test- concatenated across folds | Control | 15 min | 30 min | 1 hr | 2 hr | 4 hr | 8 hr | 24 hr | Mean | Control | 15 min | 30 min | 1 hr | 2 hr | 4 hr | 8 hr | 24 hr | Mean |
| P2=30 | 0.404 | 0.446 | 0.273 | 0.275 | 0.85 | 0.85 | 0.84 | 0.84 | 0.84 | 0.84 | 0.84 | 0.84 | 0.84 | 0.532 | 0.549 | 0.531 | 0.483 | 0.517 | 0.530 | 0.491 | 0.482 | 0.514 |
| P2=35 | 0.405 | 0.447 | 0.274 | 0.276 | 0.85 | 0.85 | 0.84 | 0.83 | 0.83 | 0.84 | 0.84 | 0.83 | 0.84 | 0.531 | 0.549 | 0.530 | 0.483 | 0.517 | 0.530 | 0.489 | 0.482 | 0.514 |
| P2=40 | 0.403 | 0.446 | 0.273 | 0.274 | 0.85 | 0.85 | 0.84 | 0.84 | 0.84 | 0.84 | 0.84 | 0.84 | 0.84 | 0.532 | 0.550 | 0.531 | 0.483 | 0.518 | 0.530 | 0.490 | 0.483 | 0.515 |
| P2=45 | 0.404 | 0.447 | 0.275 | 0.276 | 0.85 | 0.85 | 0.84 | 0.83 | 0.84 | 0.84 | 0.84 | 0.84 | 0.84 | 0.532 | 0.549 | 0.531 | 0.482 | 0.518 | 0.529 | 0.489 | 0.483 | 0.514 |
| P2=50 | 0.404 | 0.446 | 0.275 | 0.276 | 0.85 | 0.85 | 0.84 | 0.84 | 0.84 | 0.84 | 0.84 | 0.83 | 0.84 | 0.532 | 0.548 | 0.531 | 0.483 | 0.518 | 0.530 | 0.489 | 0.483 | 0.514 |
| P2=55 | 0.404 | 0.445 | 0.273 | 0.275 | 0.85 | 0.85 | 0.84 | 0.84 | 0.84 | 0.84 | 0.84 | 0.84 | 0.84 | 0.531 | 0.549 | 0.531 | 0.483 | 0.517 | 0.530 | 0.491 | 0.482 | 0.514 |
| P2=60 | 0.405 | 0.447 | 0.274 | 0.276 | 0.85 | 0.85 | 0.84 | 0.84 | 0.83 | 0.84 | 0.83 | 0.83 | 0.84 | 0.531 | 0.548 | 0.531 | 0.482 | 0.517 | 0.530 | 0.489 | 0.482 | 0.514 |
| P2=75 | 0.404 | 0.446 | 0.274 | 0.276 | 0.85 | 0.85 | 0.84 | 0.83 | 0.83 | 0.84 | 0.84 | 0.83 | 0.84 | 0.532 | 0.548 | 0.531 | 0.482 | 0.517 | 0.530 | 0.489 | 0.481 | 0.514 |
| P2=100 | 0.403 | 0.446 | 0.273 | 0.275 | 0.85 | 0.85 | 0.84 | 0.84 | 0.83 | 0.84 | 0.84 | 0.83 | 0.84 | 0.531 | 0.549 | 0.531 | 0.482 | 0.517 | 0.530 | 0.489 | 0.483 | 0.514 |

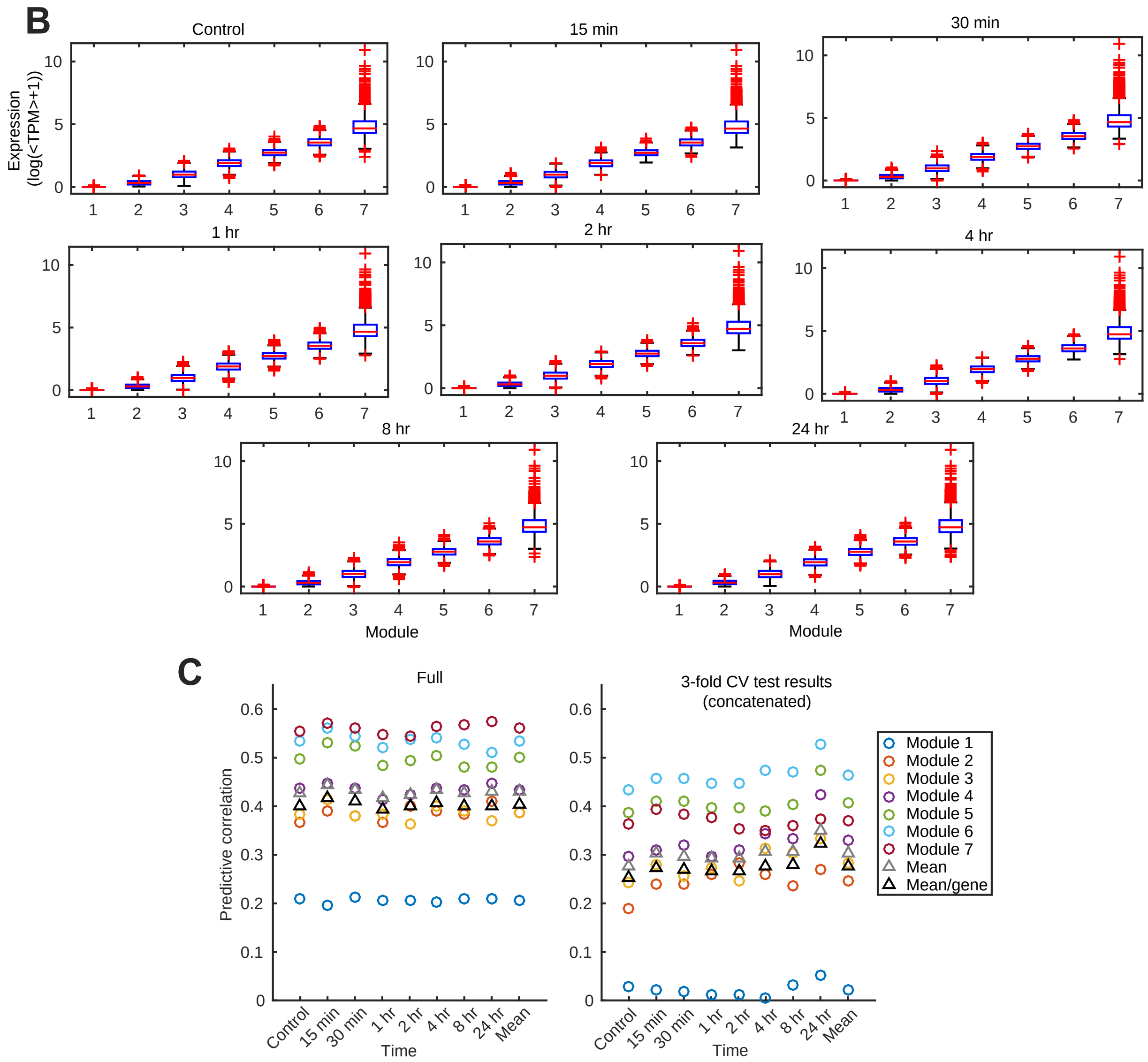


**Figure S7. DRMN hyper-parameter tuning. (A)** Summary of mean predictive power (averaged across modules) for tuning the fused-lasso hyper parameter $p_2$=30-60, given $p_1$=5 and $p_3$=0. The selected hyper-parameter setting, $p_1$=5, $p_2$=45, $p_3$=0 have the highest correlation and predictive power, justifying their selection for DRMN application. On the right are the mean similarity of

ESCAROLE and DRMN modules, and DRMN module silhouette index. **(B)** Expression levels across DRMN modules for all timepoints, presenting the significance of variation in expression across the modules, and high similarity of expression levels of the same module across time points. **(C)** Pearson correlation for the full data set (left) and for three-fold cross validation results (right). The average correlation across modules (gray triangle) and weighted average (weight corresponding to gene set size, black triangle) summarize the correlations at each time point and averaged across all time points (right-most column of points in both panels).

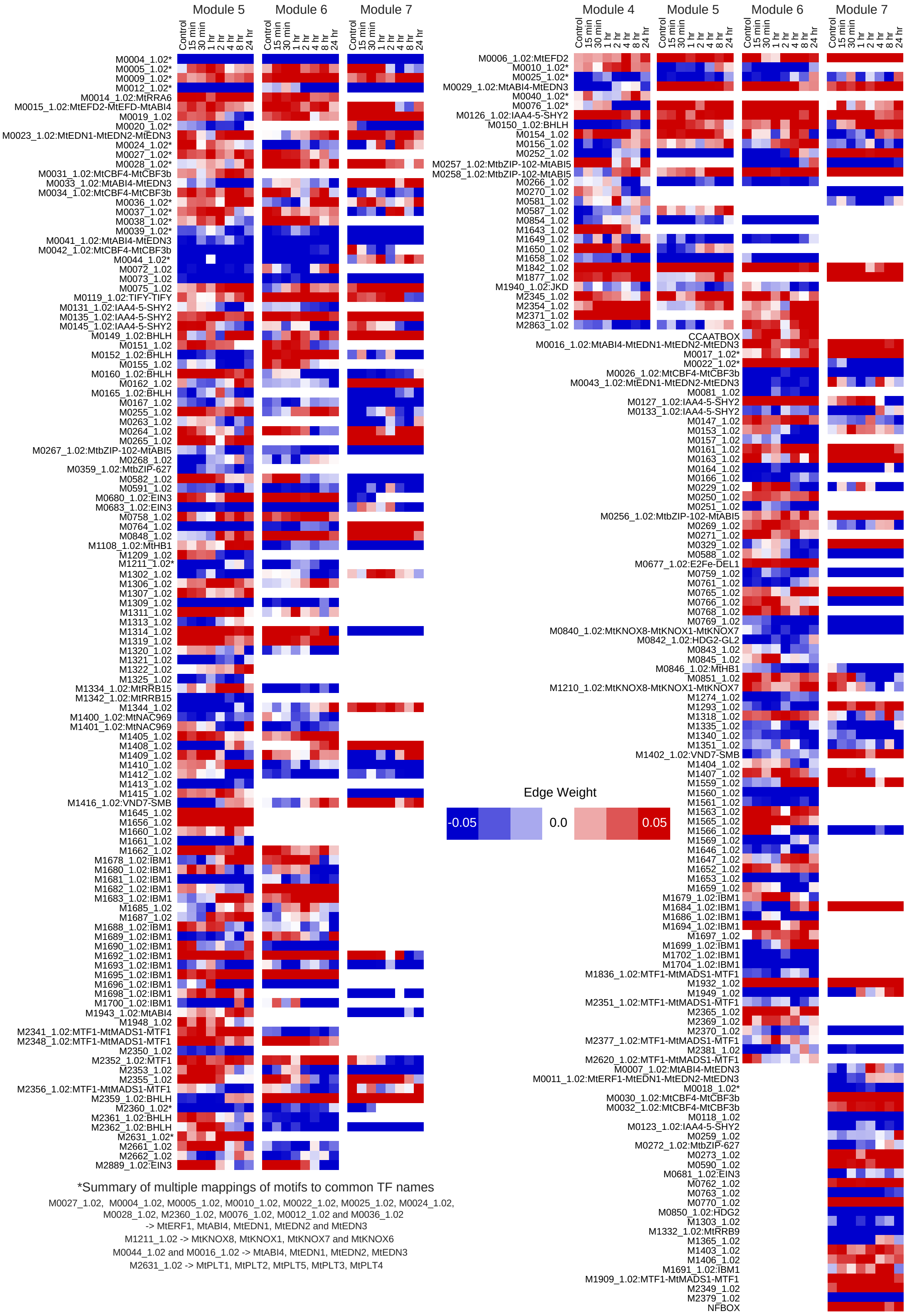


*Summary of multiple mappings of motifs to common TF names

M0027_1.02, M0004_1.02, M0005_1.02, M0010_1.02, M0022_1.02, M0025_1.02, M0024_1.02, M0028_1.02, M2360_1.02, M0076_1.02, M0012_1.02 and M0036_1.02 -> MtERF1, MtABI4, MtEDN1, MtEDN2 and MtEDN3

M1211_1.02 -> MtKNOX8, MtKNOX1, MtKNOX7 and MtKNOX6

M0044_1.02 and M0016_1.02 -> MtABI4, MtEDN1, MtEDN2, MtEDN3

M2631_1.02 -> MtPLT1, MtPLT2, MtPLT5, MtPLT3, MtPLT4

**Figure S8. DRMN edge weights for each module.** Shown are the inferred module network edge weights from the DRMN analysis. Only edges with an absolute value >0.02 in any time point and supported with enrichment (FDR-corrected hypergeometric test $P$ values < 0.05) for the corresponding motif in the respective module are shown. Edges meeting these criteria were found only for modules 4-7.

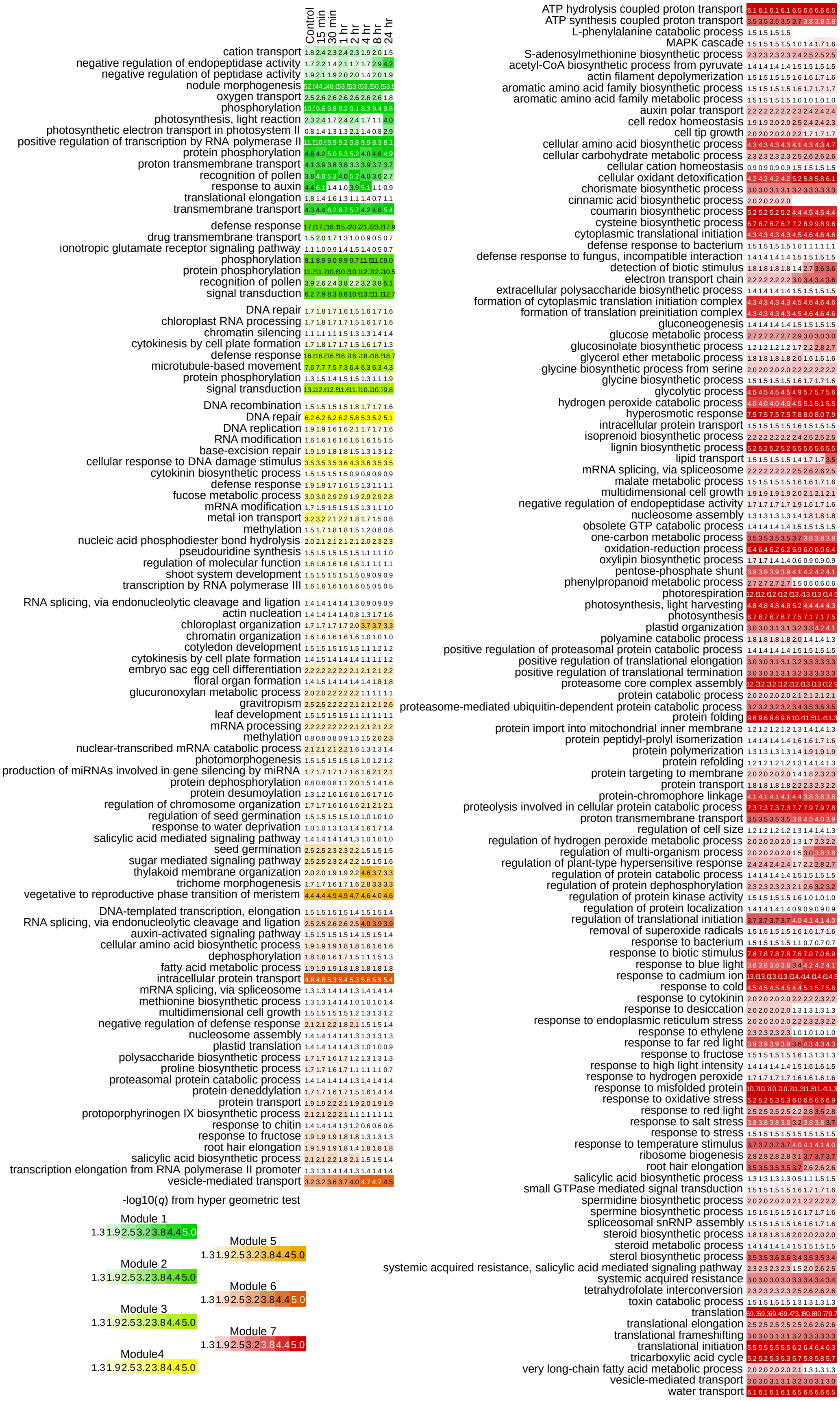
Control
15 min
30 min
1 hr
2 hr
4 hr
8 hr
24 hr
cation transport
negative regulation of endopeptidase activity
negative regulation of peptidase activity
nodule morphogenesis
oxygen transport
phosphorylation
photosynthesis, light reaction
photosynthetic electron transport in photosystem II
positive regulation of transcription by RNA polymerase II
protein phosphorylation
proton transmembrane transport
recognition of pollen
response to auxin
translational elongation
transmembrane transport
defense response
drug transmembrane transport
ionotropic glutamate receptor signaling pathway
phosphorylation
protein phosphorylation
recognition of pollen
signal transduction
DNA repair
chloroplast RNA processing
chromatin silencing
cytokinesis by cell plate formation
defense response
microtubule-based movement
protein phosphorylation
signal transduction
DNA recombination
DNA repair
DNA replication
RNA modification
base-excision repair
cellular response to DNA damage stimulus
cytokinin biosynthetic process
defense response
fucose metabolic process
mRNA modification
metal ion transport
methylation
nucleic acid phosphodiester bond hydrolysis
pseudouridine synthesis
regulation of molecular function
shoot system development
transcription by RNA polymerase II
RNA splicing, via endonucleolytic cleavage and ligation
actin nucleation
chloroplast organization
chromatin organization
cotyledon development
cytokinesis by cell plate formation
embryo sac egg cell differentiation
floral organ formation
glucuronoxylan metabolic process
gravitropism
leaf development
mRNA processing
methylation
nuclear-transcribed mRNA catabolic process
photomorphogenesis
production of miRNAs involved in gene silencing by miRNA
protein dephosphorylation
protein desumoylation
regulation of chromosome organization
regulation of seed germination
response to water deprivation
salicylic acid mediated signaling pathway
seed germination
sugar mediated signaling pathway
thylakoid membrane organization
trichome morphogenesis
vegetative to reproductive phase transition of meristem
DNA-templated transcription, elongation
RNA splicing, via endonucleolytic cleavage and ligation
auxin-activated signaling pathway
cellular amino acid biosynthetic process
dephosphorylation
fatty acid metabolic process
intracellular protein transport
mRNA splicing, via spliceosome
methionine biosynthetic process
multidimensional cell growth
negative regulation of defense response
nucleosome assembly
plastid translation
polysaccharide biosynthetic process
proline biosynthetic process
proteasomal protein catabolic process
protein deneddylation
protein transport
protoporphyrinogen IX biosynthetic process
response to chitin
response to fructose
root hair elongation
salicylic acid biosynthetic process
transcription elongation from RNA polymerase II promoter
vesicle-mediated transport
-log10(q) from hyper geometric test
Module 1
1.3 1.9 2.5 3.2 3.8 4.4 5.0
Module 2
1.3 1.9 2.5 3.2 3.8 4.4 5.0
Module 3
1.3 1.9 2.5 3.2 3.8 4.4 5.0
Module4
1.3 1.9 2.5 3.2 3.8 4.4 5.0
Module 5
1.3 1.9 2.5 3.2 3.8 4.4 5.0
Module 6
1.3 1.9 2.5 3.2 3.8 4.4 5.0
Module 7
1.3 1.9 2.5 3.2 3.8 4.4 5.0
ATP hydrolysis coupled proton transport
ATP synthesis coupled proton transport
L-phenylalanine catabolic process
MAPK cascade
S-adenosylmethionine biosynthetic process
acetyl-CoA biosynthetic process from pyruvate
actin filament depolymerization
aromatic amino acid family biosynthetic process
aromatic amino acid family metabolic process
auxin polar transport
cell redox homeostasis
cell tip growth
cellular amino acid biosynthetic process
cellular carbohydrate metabolic process
cellular cation homeostasis
cellular oxidant detoxification
chorismate biosynthetic process
cinnamic acid biosynthetic process
coumarin biosynthetic process
cysteine biosynthetic process
cytoplasmic translational initiation
defense response to bacterium
defense response to fungus, incompatible interaction
detection of biotic stimulus
electron transport chain
extracellular polysaccharide biosynthetic process
formation of cytoplasmic translation initiation complex
formation of translation preinitiation complex
gluconeogenesis
glucose metabolic process
glucosinolate biosynthetic process
glycerol ether metabolic process
glycine biosynthetic process from serine
glycine biosynthetic process
glycolytic process
hydrogen peroxide catabolic process
hyperosmotic response
intracellular protein transport
isoprenoid biosynthetic process
lignin biosynthetic process
lipid transport
mRNA splicing, via spliceosome
malate metabolic process
multidimensional cell growth
negative regulation of endopeptidase activity
nucleosome assembly
obsolete GTP catabolic process
one-carbon metabolic process
oxidation-reduction process
oxylipin biosynthetic process
pentose-phosphate shunt
phenylpropanoid metabolic process
photorespiration
photosynthesis, light harvesting
photosynthesis
plastid organization
polyamine catabolic process
positive regulation of proteasomal protein catabolic process
positive regulation of translational elongation
positive regulation of translational termination
proteasome core complex assembly
protein catabolic process
proteasome-mediated ubiquitin-dependent protein catabolic process
protein folding
protein import into mitochondrial inner membrane
protein peptidyl-prolyl isomerization
protein polymerization
protein refolding
protein targeting to membrane
protein transport
protein-chromophore linkage
proteolysis involved in cellular protein catabolic process
proton transmembrane transport
regulation of cell size
regulation of hydrogen peroxide metabolic process
regulation of multi-organism process
regulation of plant-type hypersensitive response
regulation of protein catabolic process
regulation of protein dephosphorylation
regulation of protein kinase activity
regulation of protein localization
regulation of translational initiation
removal of superoxide radicals
response to bacterium
response to biotic stimulus
response to blue light
response to cadmium ion
response to cold
response to cytokinin
response to desiccation
response to endoplasmic reticulum stress
response to ethylene
response to far red light
response to fructose
response to high light intensity
response to hydrogen peroxide
response to misfolded protein
response to oxidative stress
response to red light
response to salt stress
response to stress
response to temperature stimulus
ribosome biogenesis
root hair elongation
salicylic acid biosynthetic process
small GTPase mediated signal transduction
spermidine biosynthetic process
spermine biosynthetic process
spliceosomal snRNP assembly
steroid biosynthetic process
steroid metabolic process
sterol biosynthetic process
systemic acquired resistance, salicylic acid mediated signaling pathway
systemic acquired resistance
tetrahydrofolate interconversion
toxin catabolic process
translation
translational elongation
translational frameshifting
translational initiation
tricarboxylic acid cycle
very long-chain fatty acid metabolic process
vesicle-mediated transport
water transport

**Figure S9. Gene ontology enrichments for DRMN modules.** Each color represent a different module. The color map represents the -log10(*q)* value for the significance of enrichment, where the *q* value is the FDR corrected *P* value from a hypergeometric test for overlap.

**A**

| Threshold: No. missmatches | ESCAROLE No. genes clustered | ESCAROLE No. clusters | ESCAROLE No. GO enrichments | ESCAROLE No. clusters with GO enrichments | DRMN No. genes clustered | DRMN No. clusters | DRMN No. GO enrichments | DRMN No. clusters with GO enrichments |
|---|---|---|---|---|---|---|---|---|
| 0 | 9,771 | 228 | 72 | 33 | 8,837 | 163 | 414 | 74 |
| 1 | 10,993 | 198 | 87 | 33 | 9,633 | 133 | 462 | 64 |
| 2 | 11,612 | 112 | 80 | 31 | 10,176 | 79 | 505 | 51 |
| 3 | 11,932 | 71 | 59 | 26 | 10,409 | 55 | 505 | 39 |

**B**

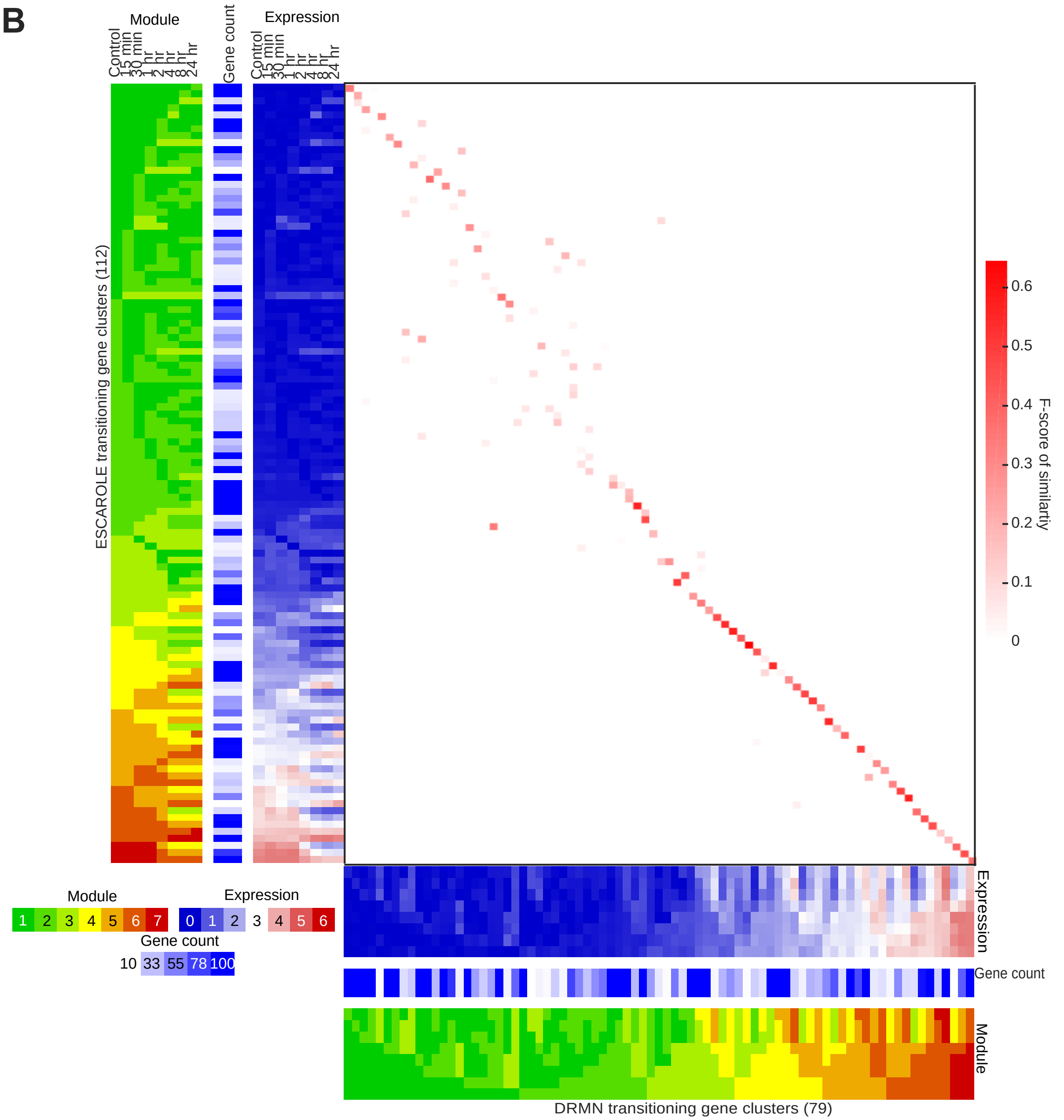


**Figure S10. Comparison of ESCAROLE and DRMN transitioning gene sets. (A)** Clustering statistics for ESCAROLE and DRMN transitioning gene sets (including numbers of gene ontology

(GO) enrichments and numbers of clusters with such enrichments) are tabulated (columns) versus the threshold for cutting the dendrogram of hierarchical clustering ( row). The thresholds represent 0-3 mismatches in module assignment across the 8-point time course (see **Methods**). The results presented in this work corresponds to 2 mismatches (purple) which includes close to maximum number of genes included, and a near-maximal number of clusters with enrichments. **(B)** Comparison of individual ESCAROLE and DRMN transitioning gene sets for gene membership. Along the vertical and horizontal axes are the mean module assignment, cluster size, and mean gene expression profiles (See also **Fig. 2B** and **Fig. 5C)**, respectively. In the center F-scores of overlaps between individual gene sets are presented (white-red color scale). The gene sets are 77% similar in gene content, but 26% similar on average for each pair of sets.

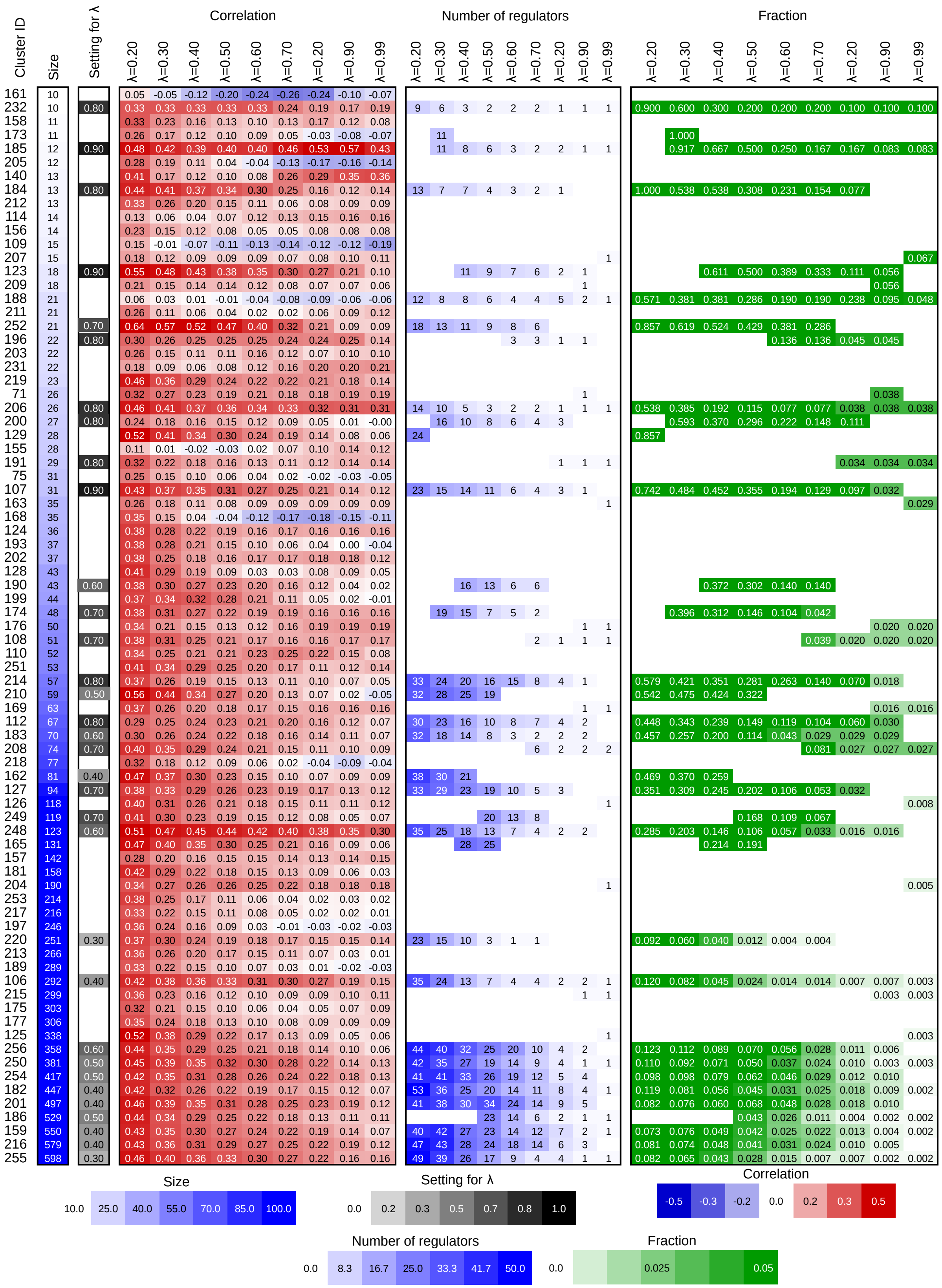

Cluster ID
Size
Setting for λ
Correlation
Number of regulators
Fraction
λ=0.20 λ=0.30 λ=0.40 λ=0.50 λ=0.60 λ=0.70 λ=0.20 λ=0.90 λ=0.99
Size
10.0 25.0 40.0 55.0 70.0 85.0 100.0
Setting for λ
0.0 0.2 0.3 0.5 0.7 0.8 1.0
Correlation
-0.5 -0.3 -0.2 0.0 0.2 0.3 0.5
Number of regulators
0.0 8.3 16.7 25.0 33.3 41.7 50.0
Fraction
0.0 0.025 0.05

**Figure S11. Hyper-parameter (λ) tuning for MTG-LASSO.** The correlation between true and predicted expression using the inferred MTG-LASSO regulator set (red heatmap), the number of significant regulators (blue heatmap center) and the ratio of significant regulators to the numbers of target genes (green) were used to choose an appropriate setting of λ for each transitioning gene cluster, indicated at the left.

**A**

*skl/ein2* mutant vs. WT DE gene sets by time point

| Larrainzar et al. experiment time-point | | Control | 30 min | 1 hr | 3 hr | 6 hr | 12 hr | 24 hr | 36 hr | 48 hr |
|---|---|---|---|---|---|---|---|---|---|---|
| DRMN module-level+motif enrichment targets (19,443) | | | | | | | | | | |
| No. overlapping | Up | 61 | 377 | 323 | 78 | 296 | 611 | 1,284 | 2,848 | 3,100 |
| | All | 92 | 596 | 466 | 149 | 442 | 1,119 | 2,388 | 5,399 | 5,638 |
| | Down | 31 | 219 | 143 | 71 | 146 | 508 | 1,104 | 2,551 | 2,538 |
| Fold enrichment | Up | 1.56 | 1.43 | 1.48 | 1.23 | 1.47 | 1.46 | 1.53 | 1.48 | 1.47 |
| | All | 1.26 | 1.27 | 1.26 | 1 | 1.18 | 1.28 | 1.33 | 1.39 | 1.36 |
| | Down | 0.92 | 1.06 | 0.93 | 0.83 | 0.84 | 1.12 | 1.16 | 1.31 | 1.24 |
| Hypergeometric *P*-value | Up | 1.2E-07 | 5.2E-25 | 8.7E-27 | 5.3E-03 | 4.3E-23 | 4.8E-45 | 1.2E-120 | 3.6E-235 | 5.5E-253 |
| | All | 7.2E-04 | 1.3E-17 | 2.8E-13 | 5.4E-01 | 2.2E-07 | 9.9E-36 | 2.2E-104 | 0.0E+00 | 2.31e-317 |
| | Down | 8.0E-01 | 1.2E-01 | 9.0E-01 | 9.9E-01 | 1.0E+00 | 1.1E-04 | 6.2E-14 | 1.7E-98 | 2.4E-63 |
| -Log10(*P*) | Up | 6.91 | 24.28 | 26.06 | 2.28 | 22.37 | 44.32 | 119.92 | 234.45 | 252.26 |
| | All | 3.14 | 16.90 | 12.56 | 0.27 | 6.67 | 35.00 | 103.66 | 300+ | 316.64 |
| | Down | 0.10 | 0.94 | 0.04 | 0.00 | 0.00 | 3.95 | 13.21 | 97.77 | 62.62 |
| MTG-LASSO predicted targets (1,607) | | | | | | | | | | |
| No. overlapping | Up | 8 | 51 | 50 | 8 | 57 | 96 | 119 | 275 | 262 |
| | All | 11 | 63 | 62 | 19 | 77 | 125 | 174 | 369 | 379 |
| | Down | 3 | 12 | 12 | 11 | 20 | 29 | 55 | 94 | 117 |
| Fold enrichment | Up | 2.48 | 2.33 | 2.78 | 1.52 | 3.41 | 2.77 | 1.71 | 1.73 | 1.5 |
| | All | 1.83 | 1.62 | 2.02 | 1.54 | 2.49 | 1.73 | 1.18 | 1.15 | 1.1 |
| | Down | 1.07 | 0.7 | 0.95 | 1.55 | 1.4 | 0.77 | 0.7 | 0.58 | 0.69 |
| Hypergeometric *P*-value | Up | 1.5E-02 | 2.3E-08 | 8.5E-11 | 1.6E-01 | 5.9E-16 | 2.1E-19 | 8.4E-09 | 6.9E-20 | 6.9E-12 |
| | All | 4.0E-02 | 1.4E-04 | 1.5E-07 | 4.3E-02 | 2.7E-13 | 1.7E-09 | 1.4E-02 | 1.3E-03 | 1.5E-02 |
| | Down | 5.3E-01 | 9.2E-01 | 6.2E-01 | 1.0E-01 | 8.3E-02 | 9.4E-01 | 1.0E+00 | 1.0E+00 | 1.0E+00 |
| -Log10(*P*) | Up | 1.82 | 7.64 | 10.07 | 0.81 | 15.23 | 18.69 | 8.07 | 19.16 | 11.16 |
| | All | 1.40 | 3.86 | 6.81 | 1.37 | 12.57 | 8.77 | 1.86 | 2.90 | 1.83 |
| | Down | 0.27 | 0.03 | 0.21 | 1.00 | 1.08 | 0.03 | 0.00 | 0.00 | 0.00 |

Larrainzar et al. experiment time-point

**B**

| Larrainzar et al. experiment time-point | Control | 30 min | 1 hr | 3 hr | 6 hr | 12 hr | 24 hr | 36 hr | 48 hr |
|---|---|---|---|---|---|---|---|---|---|
| Ranking of MTG-LASSO EIN3 targets relative to all genes by DESeq -log10(*P*) score | | | | | | | | | |
| Wilcoxon rank test | 1.50E-17 | 2.16E-64 | 6.75E-47 | 1.41E-15 | 8.93E-72 | 4.29E-76 | 2.71E-43 | 1.34E-63 | 2.57E-59 |
| GSEA (default weighted analysis) | 1.90E-01 | 1.26E-02 | 3.66E-02 | 3.95E-01 | 3.64E-02 | 1.28E-02 | 8.80E-02 | 5.32E-02 | 2.84E-01 |
| GSEA (classic mode analysis) | 7.06E-03 | 0.00E+00 | 0.00E+00 | 0.00E+00 | 0.00E+00 | 0.00E+00 | 0.00E+00 | 0.00E+00 | 0.00E+00 |

**Figure S12. Detailed statistics for EIN3 target prediction validation. A**) The putative EIN3 targets from DRMN module motif enrichment (19,443) and MTG-LASSO (1607) were overlapped with DE genes from the *skl/ein2* mutant condition relative to WT for each time point from Larrainzar et al. Shown are the numbers of predicted target genes that are also differentially expressed (yellow), fold enrichment (blue) and hypergeometric test results (*P-value*<0.05 red, otherwise gray). Both *P-value* and -log10(*P-value*) from hypergeometric tests are shown. Notably the overlaps are found to be predominantly associated with upregulated genes in the *skl/ein2* mutant condition (relative to WT). **B**) Summary of ranking results of MTG-LASSO predicted EIN3 target genes relative to all genes by DESeq -log10(*P-value*) scores of all genes for the same *skl/ein2* data (relative to WT). Shown are *P*-values of significance (*P*<0.05 red, otherwise gray) for the predicted targets to be more differentially expressed (highly ranked) than other genes in that condition from both the Wilcoxon rank test, and the GSEA "pre ranked" algorithm (in two modes; see **Methods**). Results are shown for all time points of *skl/ein2* time course.

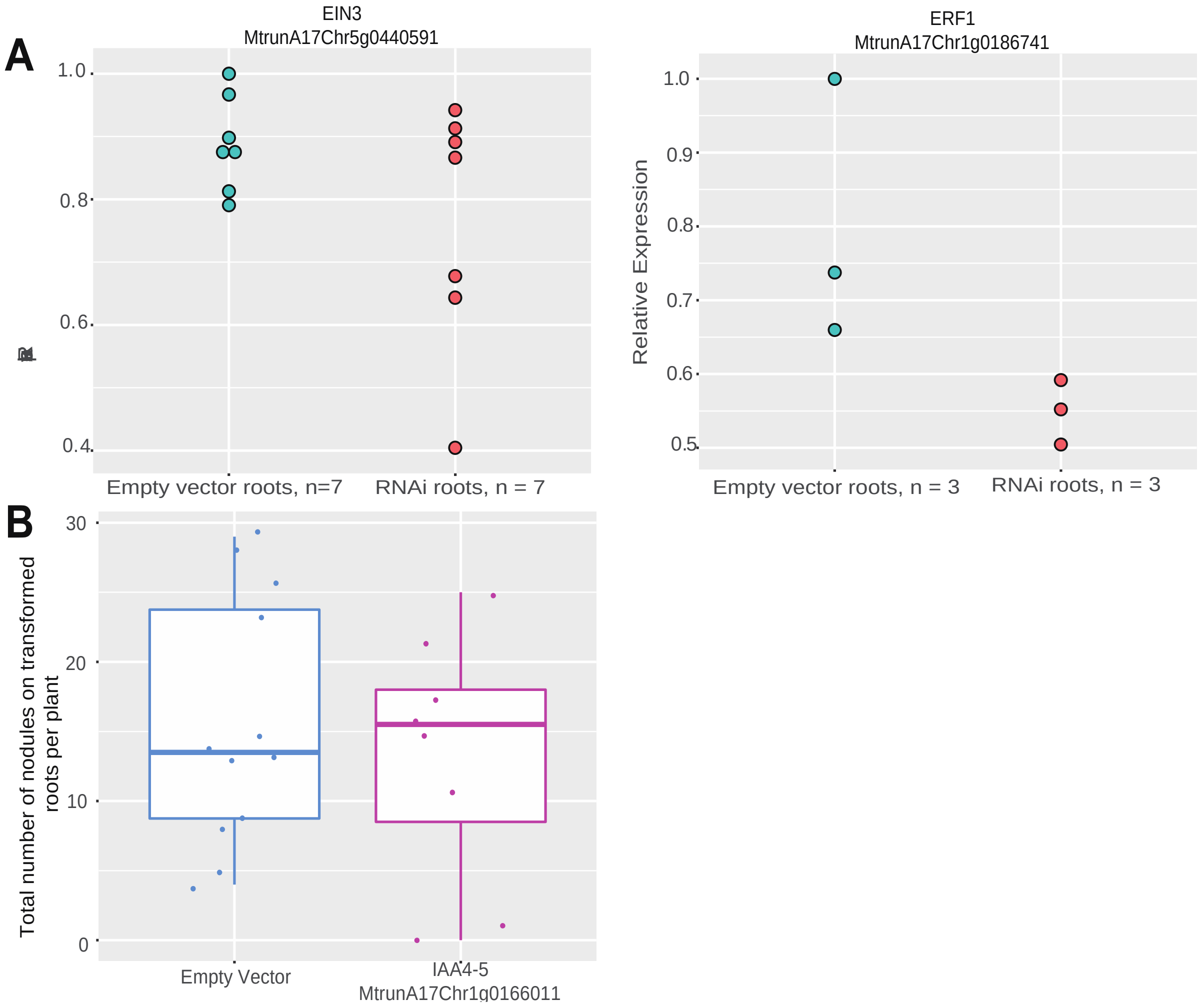


**Figure S13.** (**A**) Expression levels of MtrunA175g0440591 (EIN3) and MtrunA17Chr1g0185741 (ERF1) under RNAi knockdown with an empty vector or the respective RNAi constructs. (**B**) Knockdown of *MtrunA17Chr1g0166011* (IAA4-5) did not affect the number of nodules formed on composite *M. truncatula* plants. Radicles of Jemalong A17 were inoculated (MPMI) (26) with Agrobacterium rhizogenes MSU440, expressing the RNAi construct, or the empty vector. Three weeks after transformation, plants with transformed roots were transferred to growth pouches and inoculated with *S. meliloti* 1021 harboring pXLGD4 constitutively expressing lacZ. Two weeks after inoculation, live seedlings were stained with X-gal, and the nodules were scores and imaged.
